# Single-molecule imaging of microRNA-mediated gene silencing in cells

**DOI:** 10.1101/2021.04.30.442050

**Authors:** Hotaka Kobayashi, Robert H. Singer

**Affiliations:** Department of Anatomy and Structural Biology, Albert Einstein College of Medicine, Bronx, NY 10461, USA; PRESTO, Japan Science and Technology Agency, Chiyoda-ku, Tokyo 102-0076, Japan

## Abstract

MicroRNAs (miRNAs) are small non-coding RNAs, which regulate the expression of thousands of genes; miRNAs silence gene expression from complementary mRNAs through translational repression and mRNA decay. For decades, the function of miRNAs has been studied primarily by ensemble methods, where a bulk collection of molecules is measured outside cells. Thus, the behavior of individual molecules during miRNA-mediated gene silencing, as well as their spatiotemporal regulation inside cells, remains mostly unknown. Here we report single-molecule methods to visualize each step of miRNA-mediated gene silencing *in situ* inside cells. Simultaneous visualization of single mRNAs, translation, and miRNA-binding revealed that miRNAs preferentially bind to translated mRNAs rather than untranslated mRNAs. Spatiotemporal analysis based on our methods uncovered that miRNAs bind to mRNAs immediately after nuclear export. Subsequently, miRNAs induced translational repression and mRNA decay within 30 and 60 min, respectively, after the binding to mRNAs. This methodology provides a framework for studying mRNA regulation at the single-molecule level with spatiotemporal information inside cells.

## Introduction

MicroRNAs (miRNAs) are ∼22-nt small non-coding RNAs, which silence gene expression from complementary mRNAs **(*1*–*5*)**. Within the human genome, there are over 2000 miRNAs **(*6*)**, which regulate the expression of thousands of mRNAs **(*7*)**, thereby influencing various biological processes and diseases. Notably, miRNAs cannot work alone; they assemble with the Argonaute subfamily of proteins (AGO) into the effector complex called the RNA-induced silencing complex (RISC) **(*8*–*10*)**. Using miRNAs as guides, RISC binds to the 3′ UTR of target mRNAs **(*11*–*13*)**, inducing translational repression, followed by mRNA decay **(*14*–*18*)**.

After the discovery of the first miRNA in 1993 **(*1, 2*)**, miRNA-mediated gene silencing has been studied for decades. However, the function of miRNAs has been monitored primarily by ensemble methods, e.g., luciferase assays and RNA sequencing, where a bulk collection of molecules is measured outside cells **(*8*–*10, 14*–*18*)**. Thus, the behavior of individual molecules during miRNA-mediated gene silencing, as well as their spatiotemporal regulation inside cells, remains mostly unknown. Here we report a series of single-molecule methods to visualize each step of miRNA-mediated gene silencing, i.e., RISC-binding, translational repression, and mRNA decay, *in situ* inside cells. As our methods visualize the function of miRNAs on a cell-by-cell basis, they enable both single-molecule and single-cell analysis. These technical advantages, which overcome the limitation of canonical methods, provide novel insights into when, where, and how miRNAs work inside cells.

## Results

### Visualization of mRNA decay by miRNAs with single-molecule resolution

First, we sought to develop a method to visualize miRNA-mediated mRNA decay with single-molecule resolution. In human U2OS cells, which have been widely used for RNA imaging **(*19*–*21*)**, miR-21 is the most abundant miRNA **(*22*)** (fig. S1, a to c). Thus, we constructed the reporter system that recapitulates mRNA decay by miR-21 (Fig. 1a). In this reporter system, where two different mRNAs are expressed under the control of a bi-directional promoter, *firefly luciferase* (*Fluc*) mRNAs represent the internal control. *SunTag* mRNAs with miR-21 sites are used to monitor miRNA-mediated mRNA decay, while *SunTag* mRNAs with mutant sites are used as the negative control (fig. S1d). In this method, reporter mRNAs are detected by single-molecule fluorescence *in situ* hybridization (smFISH) to visualize them with single-molecule sensitivity **(*23, 24*)** (Fig. 1b, and fig. S2).

**Fig. 1.**
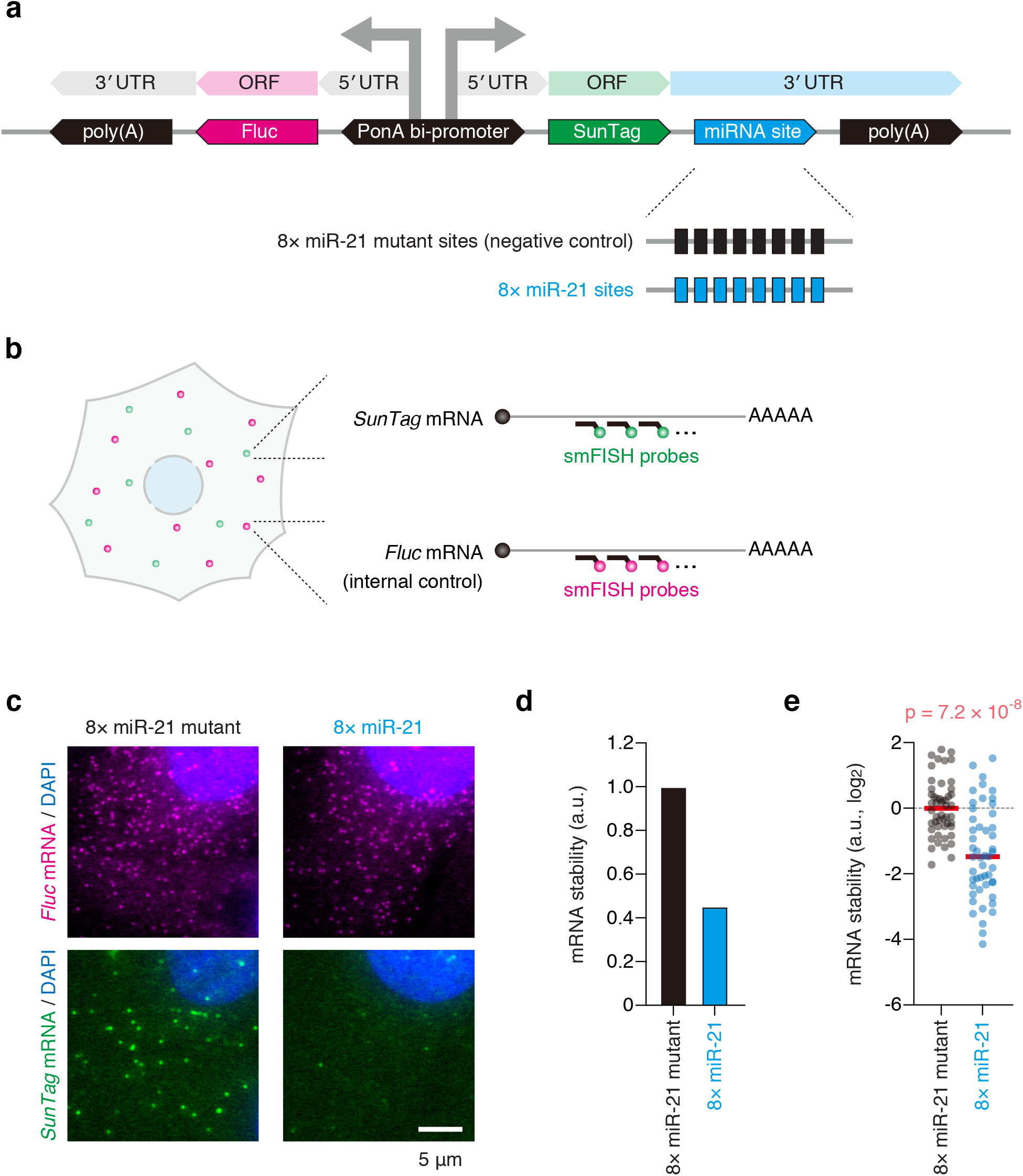
Single-molecule imaging of miRNA-mediated mRNA decay. **(a)** Schematic of the reporter construct to recapitulate miRNA-mediated mRNA decay. PonA bi-promoter, PonA-inducible bi-directional promoter. **(b)** Schematic of the smFISH experiment to visualize miRNA-mediated mRNA decay at single-mRNA resolution. Green and magenta spots represent *SunTag* and *Fluc* mRNAs, respectively. **(c)** The images of U2OS cells expressing the 8× miR-21 mutant reporter (left) and the 8× miR-21 reporter (right). *Fluc* mRNAs (magenta, top) and *SunTag* mRNAs (green, bottom) were labeled by smFISH. Nuclei (blue) were stained by DAPI. Scale bar, 5μm. (**d** and **e**) mRNA decay mediated by miR-21. Images were analyzed using CellProfiler and FISH-quant. Then, mRNA stability was calculated as described in fig. S2 (see also Online Methods). The results of bulk analysis (d) and single-cell analysis (e) are shown. In (e), each circle represents a single cell (n = 50 for each condition), while red lines represent the medians. The p value of Mann Whitney test is shown.

Validating our method, U2OS cells expressing *SunTag* mRNAs with miR-21 sites showed a smaller number of mRNAs, compared with the negative control (Fig. 1c). For quantitative analysis, we performed three-dimensional (3D) fluorescence imaging, followed by single-molecule detection in 3D using the FISH-quant algorithm **(*25*)** (fig. S2). This analysis confirmed the reduction of mRNA stability when *SunTag* mRNAs have miR-21 sites (Fig. 1, d and e, and fig. S3, a to c). Although ensemble methods analyze a bulk collection of mRNAs from numerous cells **(*8*–*10, 14, 16*–*18, 22*)**, our method can analyze miRNA-mediated mRNA decay on a cell-by-cell basis. This advantage highlighted the cellular heterogeneity of miRNA-mediated mRNA decay (Fig. 1e). Notably, unlike canonical methods, where relative expression levels are analyzed, our method can count the absolute number of mRNAs (fig. S3, a and b), thereby allowing the absolute quantification of miRNA function.

### Visualization of translational repression by miRNAs with single-molecule resolution

Second, we attempted to develop a method to visualize miRNA-mediated translational repression with single-molecule resolution. To this end, we took advantage of the technique called single-molecule imaging of nascent peptides (SINAPS) **(*21*)**, where translation of reporter mRNAs can be visualized with single-molecule resolution. Based on the principle of SINAPS, we constructed the reporter mRNA that recapitulates translational repression by miR-21 (Fig. 2a). This reporter has the SunTag sequence, consisting of 24 tandem repeats of the GCN4 epitope **(*26*)**, in the ORF. Through immunofluorescence (IF) with anti-GCN4 antibodies, SunTag allows us to visualize nascent peptides being translated from mRNAs with single-molecule sensitivity. To inhibit the accumulation of SunTag throughout the cytoplasm, which dramatically increases background fluorescence, the degron sequence was added to the C terminus of the ORF **(*27*)**. To monitor miRNA-mediated translational repression, miR-21 sites were inserted in the 3′ UTR. As miRNAs also trigger mRNA decay (Fig. 1c), which causes a non-negligible reduction in the number of mRNAs for analysis, we added the anti-decay sequence, A_114_-N_40_ **(*28, 29*)**, to the end of the 3′ UTR. In this method, reporter mRNAs and their translation are visualized by smFISH and IF, respectively, with single-molecule resolution (Fig. 2b, and fig. S4).

**Fig. 2.**
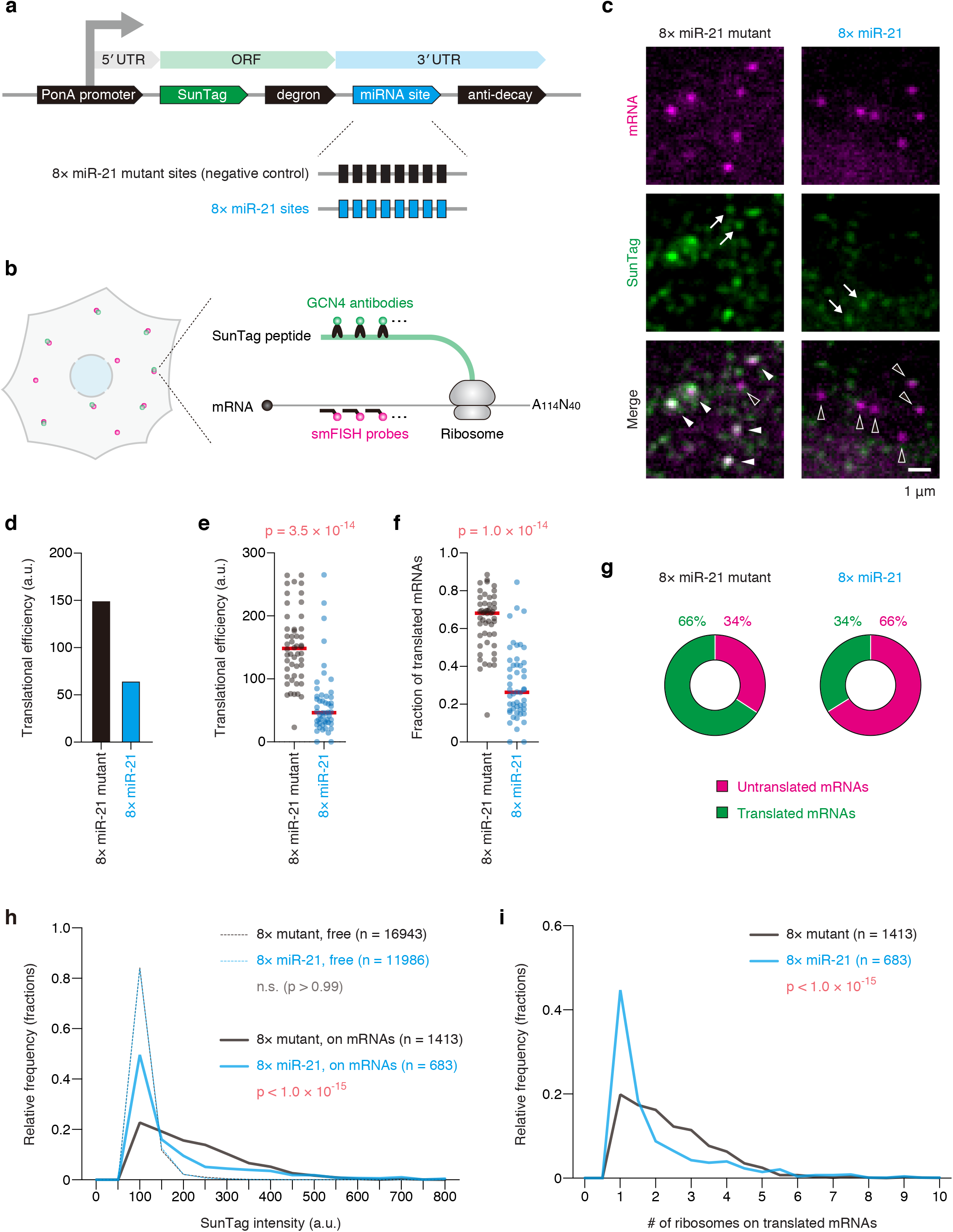
Single-molecule imaging of miRNA-mediated translational repression. **(a)** Schematic of the reporter construct to recapitulate miRNA-mediated translational repression. PonA promoter, PonA-inducible promoter. **(b)** Schematic of the SINAPS experiment to visualize miRNA-mediated translational repression at single-mRNA resolution. Green and magenta spots represent SunTag peptides and reporter mRNAs, respectively. **(c)** The images of U2OS cells expressing the 8× miR-21 mutant reporter (left) and the 8× miR-21 reporter (right). Reporter mRNAs (magenta, top) and SunTag peptides (green, middle) were labeled by SINAPS. Merged images are shown at the bottom. White and black arrow heads indicate translated and untranslated mRNAs, respectively, while white arrows indicate “free” SunTag peptides. Scale bar, 1μm. (**d** and **e**) Translational repression mediated by miR-21. Images were analyzed using CellProfiler and FISH-quant. Then, translational efficiency was calculated as described in fig. S4 (see also Online Methods). The results of bulk analysis (d) and single-cell analysis (e) are shown. In (e), each circle represents a single cell (n = 50 for each condition), while red lines represent the medians. The p value of Mann Whitney test is shown. **(f)** Reduction of the fraction of translated mRNAs by miR-21. The fraction of translated mRNAs was calculated as described in fig. S4 (see also Online Methods). Each circle represents a single cell (n = 50 for each condition), while red lines represent the medians. The p value of Mann Whitney test is shown. **(g)** The ratio of untranslated and translated mRNAs. All mRNAs were classified into untranslated or translated mRNAs based on 3D colocalization analysis. **(h)** The histogram of SunTag intensity. The intensities of free SunTag spots (dashed lines) and of SunTag spots on mRNAs (solid lines) are shown. The p values of Dunn’s multiple comparisons test are shown. n.s., not significant. **(i)** Reduction of the number of ribosomes on translated mRNAs by miR-21. The number of ribosomes on translated mRNAs was calculated as described in fig. S4 (see also Online Methods). The p value of Mann Whitney test is shown.

U2OS cells expressing reporter mRNAs without miR-21 sites showed bright SunTag signals on mRNAs (Fig. 2c, left panels, white arrowheads), indicating that translation is successfully visualized. In agreement with this, SunTag signals on mRNAs almost completely disappeared upon the treatment with puromycin, an inhibitor of translation (fig. S5, a to c). Importantly, U2OS cells expressing reporter mRNAs with miR-21 sites did not show bright SunTag signals on mRNAs (Fig. 2c, right panels, black arrowheads). The reduction of translational efficiency by miR-21 was confirmed by quantitative analysis (Fig. 2, d and e, and fig. S6, a and b). Together, these results indicate that our method makes it possible to visualize miRNA-mediated translational repression with single-molecule resolution. Single-cell analysis based on this method revealed the cellular heterogeneity of miRNA-mediated translational repression (Fig. 2e), the same as miRNA-mediated mRNA decay (Fig. 1e).

Canonical methods, where a bulk collection of mRNAs is analyzed, are sufficient to monitor translational repression by miRNAs **(*15*–*18, 28, 29*)**. However, even if these methods detect a 50% reduction in translational efficiency, they cannot address how miRNAs accomplished the 50% silencing inside cells; the number of translated mRNAs may be reduced to 50%, or the number of ribosomes on translated mRNAs may be reduced to 50%. Taking advantage of single-molecule methods, we next addressed this issue. To identify the number of translated mRNAs, we performed 3D colocalization analysis between mRNAs and SunTag. In this analysis, the 3D positions of mRNAs and SunTag were localized at sub-pixel resolution by 3D Gaussian fitting **(*25*)**. Subsequently, based on the colocalization with SunTag, which is determined by the 3D distance, all mRNAs were classified into “untranslated” or “translated” mRNAs (fig. S4). This analysis revealed that miRNAs reduce the number of translated mRNAs within cells (Fig. 2, f and g). Notably, translation of reporter mRNAs was completely halted by miRNAs in a few cells (Fig. 2f, see cells at the bottom). In SINAPS experiments, dim SunTag signals that do not colocalize with mRNAs (free SunTag) represent single SunTag peptides released from ribosomes **(*21*)** (Fig. 2c, arrows). On the other hand, bright SunTag signals consist of multiple SunTag peptides being translated by multiple ribosomes (Fig. 2c, white arrowheads). Thus, using the fluorescence intensity of free SunTag and that of SunTag on mRNAs (Fig. 2h), the number of ribosomes on translated mRNAs could be roughly estimated **(*21*)** (fig. S4). This analysis revealed that miRNAs also reduced the number of ribosomes on translated mRNAs (Fig. 2i).

### Imaging of RISC-binding with single-molecule resolution

Third, we sought to establish a method to image RISC-binding with single-molecule resolution. To this end, the reporter mRNA for translational repression, harboring eight miR-21 sites (Fig. 2a), was repurposed to image RISC on mRNAs. In this method, RISC was imaged by IF with anti-AGO antibodies, while reporter mRNAs were imaged by smFISH with single-molecule resolution (Fig. 3a, and fig. S7). Although there are four AGO proteins (AGO1-4) in humans **(*12*)**, AGO2 is predominantly expressed in U2OS cells **(*30*)** (fig. S8a), hence we focused on AGO2 in this study. For our method, it was crucial to eliminate the unwanted RISC-binding to reporter mRNAs independent of miR-21. Thus, we carefully removed the potential miRNA sites (8mer, 7mer, and 6mer) **(*11*)** of the top 30 most abundant miRNAs from the reporter mRNAs (fig. S1, b and c).

**Fig. 3.**
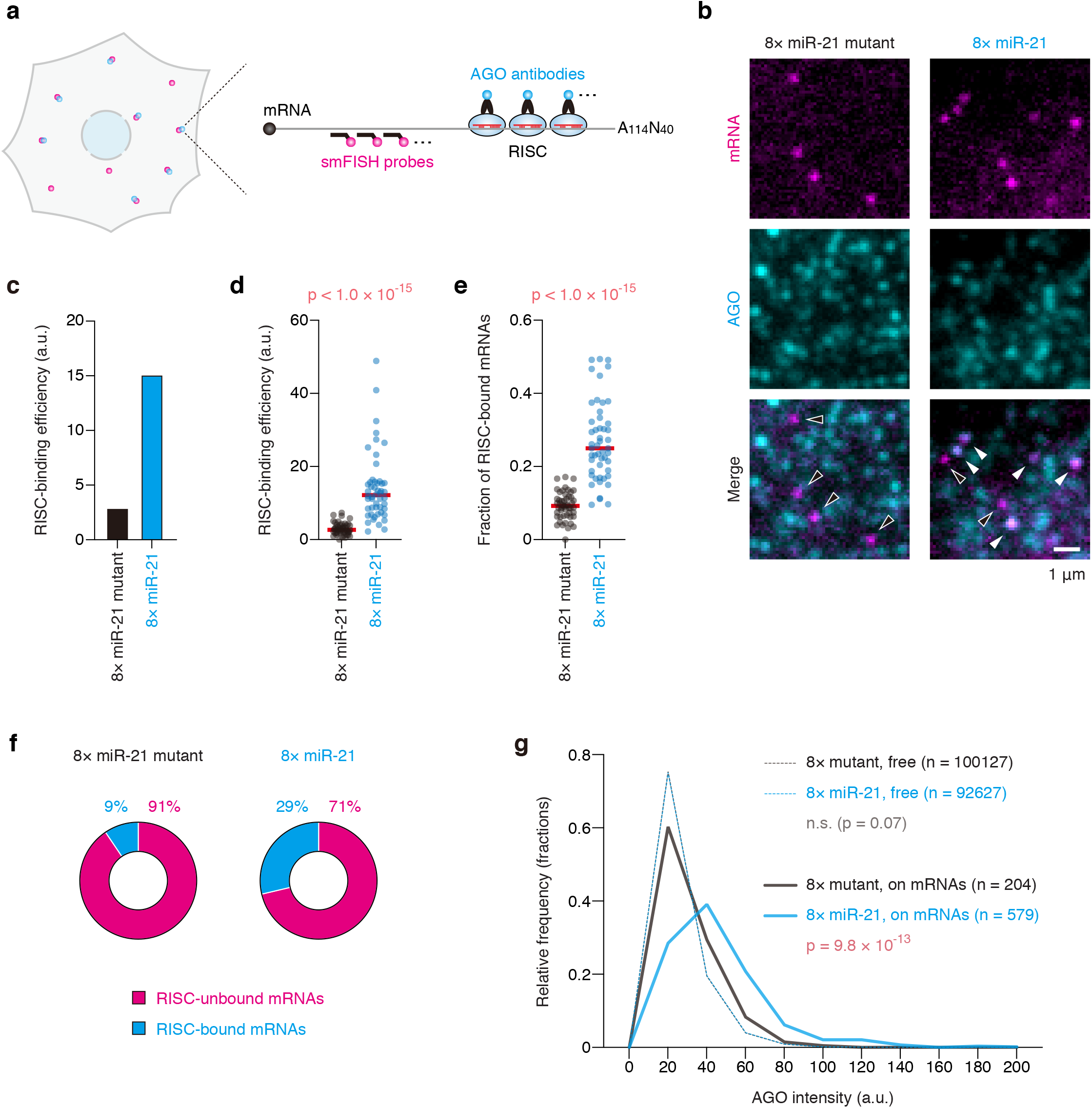
Single-molecule imaging of RISC-binding. **(a)**Schematic of the IF-FISH experiment to visualize RISC-binding at single-mRNA resolution. Cyan and magenta spots represent RISC and reporter mRNAs, respectively. **(b)**The images of U2OS cells expressing the 8× miR-21 mutant reporter (left) and the 8× miR-21 reporter (right). Reporter mRNAs (magenta, top) and AGO (cyan, middle) were labeled by IF-FISH. Merged images are shown at the bottom. White and black arrow heads indicate RISC-bound and RISC-unbound mRNAs, respectively. Scale bar, 1μm. (**c** and **d**) RISC-binding mediated by miR-21. Images were analyzed using CellProfiler and FISH-quant. Then, RISC-binding efficiency was calculated as described in fig. S7 (see also Online Methods). The results of bulk analysis (c) and single-cell analysis (d) are shown. In (d), each circle represents a single cell (n = 50 for each condition), while red lines represent the medians. The p value of Mann Whitney test is shown. **(e)** Increase of the fraction of RISC-bound mRNAs by miR-21. The fraction of RISC-bound mRNAs was calculated as described in fig. S7 (see also Online Methods). Each circle represents a single cell (n = 50 for each condition), while red lines represent the medians. The p value of Mann Whitney test is shown. **(f)** The ratio of RISC-unbound and RISC-bound mRNAs. All mRNAs were classified into RISC-unbound or RISC-bound mRNAs based on 3D colocalization analysis. **(g)** The histogram of AGO intensity. The intensities of free AGO spots (dashed lines) and of AGO spots on mRNAs (solid lines) are shown. The p values of Dunn’s multiple comparisons test are shown. n.s., not significant.

Given the number of AGO proteins (fig. S8a, ∼15,000 molecules per cell) and the relative occupancy of miR-21 (fig. S1c, ∼25% of miRNAs) in U2OS cells, the number of RISC loaded with miR-21 is estimated to be ∼4000 per cell. Thus, we minimized the expression level of reporter mRNAs (maximum, ∼100; median ∼40 mRNAs per cell), so that reporter mRNAs are efficiently recognized by RISC. Validating our method, U2OS cells expressing reporter mRNAs with miR-21 sites showed AGO signals on mRNAs (Fig. 3b, right panels, white arrowheads). In contrast, reporter mRNAs with mutant sites did not colocalize with AGO (Fig. 3b, left panels, black arrowheads). RISC-binding mediated by miR-21 was confirmed quantitatively by bulk analysis (Fig. 3c), single-cell analysis (Fig. 3, d and e, and fig. S8b), and single-molecule analysis (Fig. 3, f and g). As with mRNA decay and translational repression (Fig. 1e, and Fig. 2e), our method highlighted the cell-to-cell heterogeneity of RISC-binding efficiency (Fig. 3d).

### Simultaneous visualization of single mRNAs, translation, and RISC-binding

Since we developed a series of methods to visualize each step of miRNA-mediated gene silencing with single-molecule resolution, we next explored the relationship between these steps at the single-mRNA level. To this end, we visualized single mRNAs, translation, and RISC-binding simultaneously (Fig. 4a, and fig. S9), using the reporter mRNA harboring miR-21 sites (Fig. 2a). Based on 3D colocalization analysis, all mRNAs were classified into four classes: 1) RISC-unbound untranslated mRNAs, 2) RISC-unbound translated mRNAs, 3) RISC-bound untranslated mRNAs, and 4) RISC-bound translated mRNAs (Fig. 4, b and c). When we used these data for single-cell analysis, RISC-binding efficiency and translational efficiency were negatively correlated (Fig. 4d), validating our experiments.

**Fig. 4.**
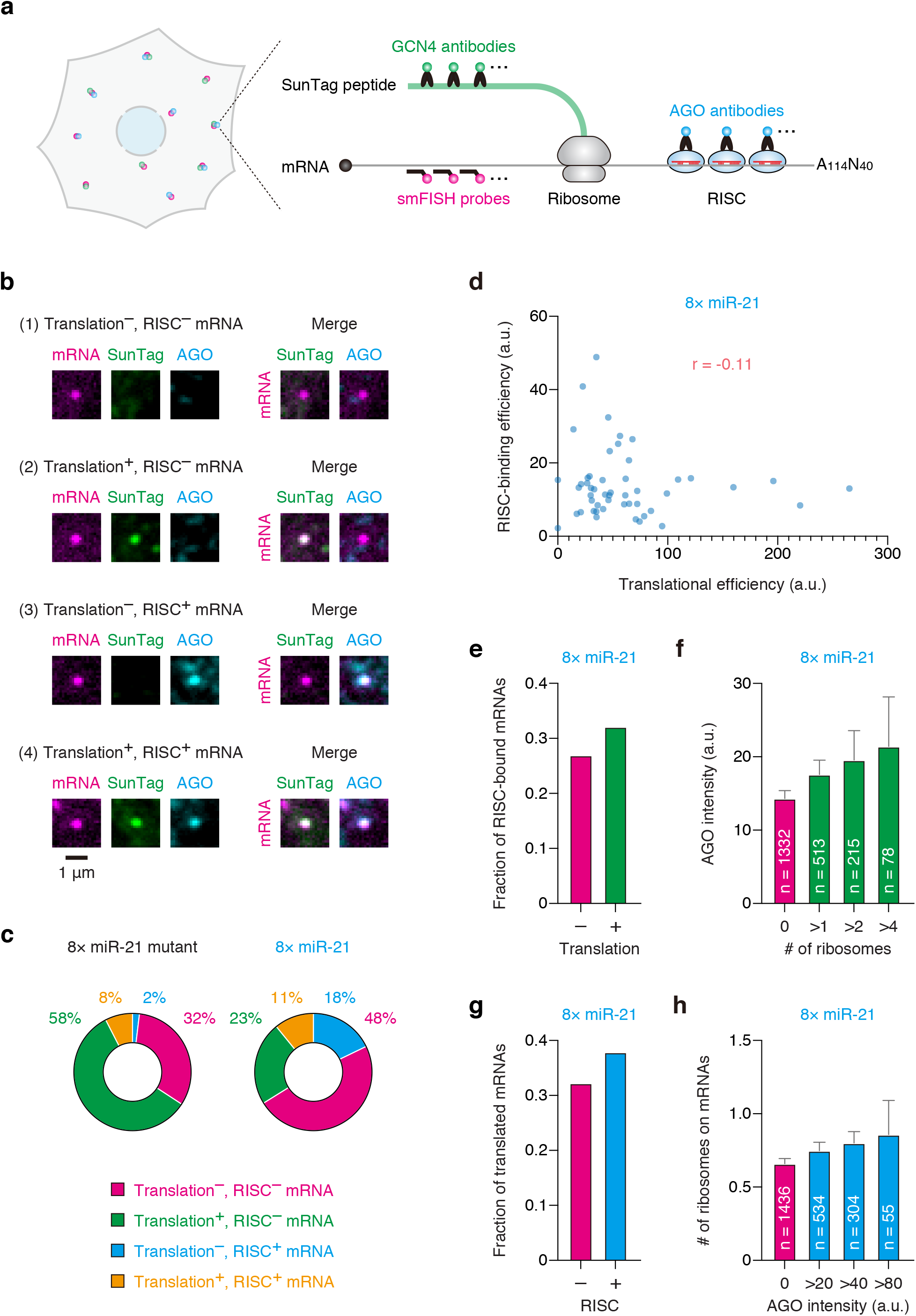
Simultaneous visualization of single mRNAs, translation, and RISC-binding. **(a)** Schematic of the SINAPS-IF-FISH experiment to visualize translation and RISC-binding simultaneously at single-mRNA resolution. Magenta, green, and cyan spots represent reporter mRNAs, SunTag peptides, and RISC, respectively. **(b)** The images of a RISC-unbound untranslated mRNA (first row), a RISC-unbound translated mRNA (second row), a RISC-bound untranslated mRNA (third row), and a RISC-bound translated mRNA (fourth row) in U2OS cells are shown. Reporter mRNAs (magenta, first column), SunTag peptides (green, second column), and AGO (cyan, third column) were labeled by SINAPS and IF-FISH. Merged images are shown on the right side. Scale bar, 1μm. **(c)** The ratio of RISC-unbound untranslated (magenta), RISC-unbound translated (green), RISC-bound untranslated (cyan), and RISC-bound translated (orange) mRNAs. All mRNAs were classified into these four classes based on 3D-colocalization analysis. **(d)** Negative correlation between translational efficiency and RISC-binding efficiency at the single-cell level. Images were analyzed using CellProfiler and FISH-quant. Then, translational efficiency and RISC-binding efficiency were calculated as described in fig. S9 (see also Online Methods). Each circle represents a single cell (n = 50). The Pearson correlation coefficient (r) is shown. (**e** and **f**) Translated mRNAs tend to be RISC-bound mRNAs. The fraction of RISC-bound mRNAs (e) and the intensity of AGO on mRNAs (f) are shown. In (f), the means with SEM are shown. Magenta and green bars represent the values of untranslated and translated mRNAs, respectively. (**g** and **h**) RISC-bound mRNAs tend to be translated mRNAs. The fraction of translated mRNAs (g) and the number of ribosomes on mRNAs (h) are shown. In (h), the means with SEM are shown. Magenta and cyan bars represent the values of RISC-unbound and RISC-bound mRNAs, respectively.

Unexpectedly, single-mRNA analysis revealed that translated mRNAs tend to be bound by RISC, compared with untranslated mRNAs (Fig. 4e). This tendency was confirmed by the quantitative analysis, where we analyzed the fluorescence intensity of AGO on mRNAs with different numbers of ribosomes (Fig. 4f). In line with this, RISC-bound mRNAs tend to be translated, compared with RISC-unbound mRNAs (Fig. 4g). The quantitative analysis confirmed that the mRNAs efficiently bound by RISC are efficiently translated (Fig. 4h). Given that RISC does not activate translation, these data indicate that RISC preferentially binds to translated mRNAs rather than untranslated mRNAs.

### Spatiotemporal analysis of RISC-binding, translational repression, and mRNA decay by single-mRNA imaging

Even though RISC prefers translated mRNAs over untranslated mRNAs, if RISC represses translation immediately, most of RISC-bound mRNAs should be untranslated mRNAs. As our data showed the opposite (Fig. 4, e to h), we speculated that RISC needs a relatively long time to repress translation. Taking advantage of our methods, which can visualize RISC-binding, translational repression, and mRNA decay, at the single-mRNA and single-cell levels, we next explored the time course of miRNA-mediated gene silencing.

Firstly, using the methods we developed (Figs. 2a and 4a), we performed spatiotemporal analysis of RISC-binding and translational repression simultaneously. In this analysis, transcription of the reporter mRNAs was strictly controlled under the control of the Ponasterone A (PonA)-inducible promoter **(*31, 32*)** (Fig. 2a); after the pulse of PonA treatment, reporter mRNAs were observed by single-mRNA imaging at three different time points (Fig. 5a). During these experiments, the outlines of the nuclei and cells, visualized by DAPI and the non-specific background signals of smFISH probes, respectively, were automatically detected by the CellProfiler algorithm **(*33*)** (fig. S9). Validating our experiments, the ratio of the number of cytoplasmic mRNAs to that of nuclear mRNAs was increased over time by nuclear export (Fig. 5b). Notably, single-cell analysis revealed that RISC binds to cytoplasmic mRNAs as early as at 0 min after the completion of PonA treatment (Fig. 5c, and fig. S10a). These cytoplasmic mRNAs should be the mRNAs immediately after nuclear export, because most mRNAs are still in the nucleus at this time point (Fig. 5b). Single-mRNA analysis, where we analyzed the intensity of AGO on cytoplasmic mRNAs, confirmed that RISC-binding takes place instantly (Fig. 5d, and fig. S10c). When we focused on translational repression, however, single-cell analysis showed that RISC does not repress translation until 30 min after the PonA pulse (Fig. 5e, and fig. S10b). This was confirmed by single-mRNA analysis, where we analyzed the number of ribosomes on translated mRNAs in the cytoplasm (Fig. 5f, and fig. S10c). Together, these results indicate that RISC binds to mRNAs immediately after their transport from the nucleus to the cytoplasm, followed by translational repression within 30 min. The data of spatiotemporal analysis also confirmed that RISC prefers translated mRNAs over untranslated mRNAs (Fig. 5, g and h).

**Fig. 5.**
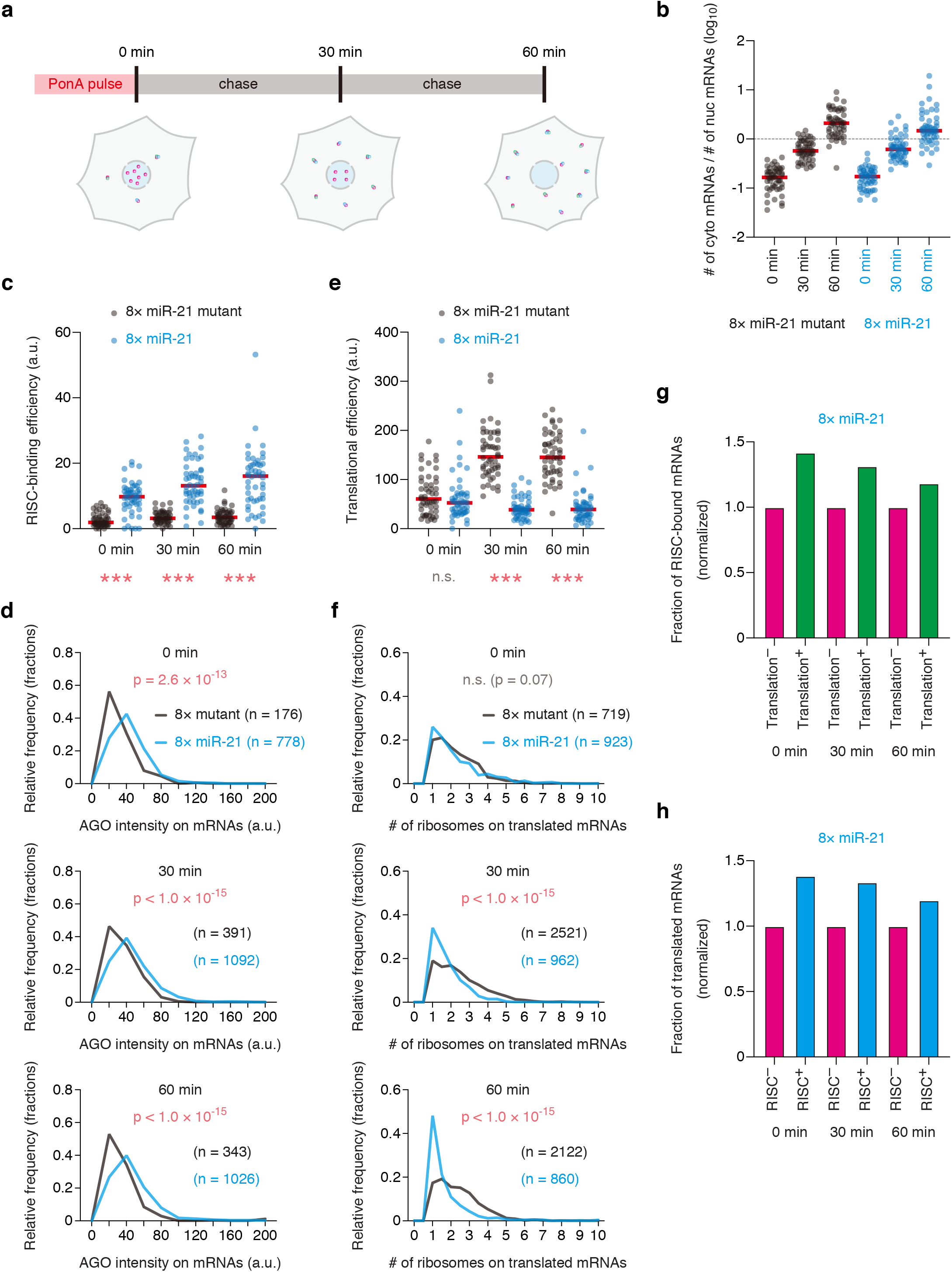
Spatiotemporal analysis of RISC-binding and translational repression by single-mRNA imaging. **(a)** Schematic of spatiotemporal analysis of RISC-binding and translational repression by single-mRNA imaging. In pulse-chase experiments, IF-FISH and SINAPS were performed to visualize RISC-binding and translation at single-mRNA resolution. Magenta, green, and cyan spots represent reporter mRNAs, SunTag peptides, and RISC, respectively. **(b)** Transport of reporter mRNAs from the nucleus to the cytoplasm. In pulse-chase experiments, reporter mRNAs were labeled by smFISH. The ratios of the number of cytoplasmic mRNAs to that of nuclear mRNAs are shown. Each circle represents a single cell (n = 50 for each condition), while red lines represent the medians. cyto, cytoplasmic. nuc, nuclear. (**c** to **f**) Time-course analysis of RISC-binding (c and d) and translational repression (e and f) by single-mRNA imaging. Images were analyzed using CellProfiler and FISH-quant. Then, RISC-binding efficiency (c), the intensity of AGO on mRNAs (d), translational efficiency (e), and the number of ribosomes on translated mRNAs (f) were calculated as described in fig. S9 (see also Online Methods). In (c) and (e), each circle represents a single cell (n = 50 for each condition), while red lines represent the medians. The p values of Dunn’s multiple comparisons test are shown. *** and n.s. represent p < 0.001 and not significant (p > 0.05), respectively. **(g)** Translated mRNAs tend to be RISC-bound mRNAs at all time points. The fraction of RISC-bound mRNAs is shown. Magenta and green bars represent the values of untranslated and translated mRNAs, respectively. **(h)** RISC-bound mRNAs tend to be translated mRNAs at all time points. The fraction of translated mRNAs is shown. Magenta and cyan bars represent the values of RISC-unbound and RISC-bound mRNAs, respectively.

Finally, we performed spatiotemporal analysis of mRNA decay using our reporter system (Fig. 1a). Under the control of the PonA-inducible bi-directional promoter, *Fluc* mRNAs, the internal control, and *SunTag* mRNAs, the reporter to monitor mRNA decay, were expressed for time-course experiments (Fig. 6a). As with the analysis for RISC-binding and translational repression (Fig. 5b), the ratio of the number of cytoplasmic mRNAs to that of nuclear mRNAs was increased over time (Fig. 6b), indicating that our spatiotemporal analysis worked well. Unlike translational repression, mRNA decay was not observed at 30 min after the PonA pulse (Fig. 6c). Instead, single-cell analysis showed a reduction of mRNA stability at 60 min after the PonA pulse, indicating that RISC induces mRNA decay within 60 min after the binding to mRNAs.

**Fig. 6.**
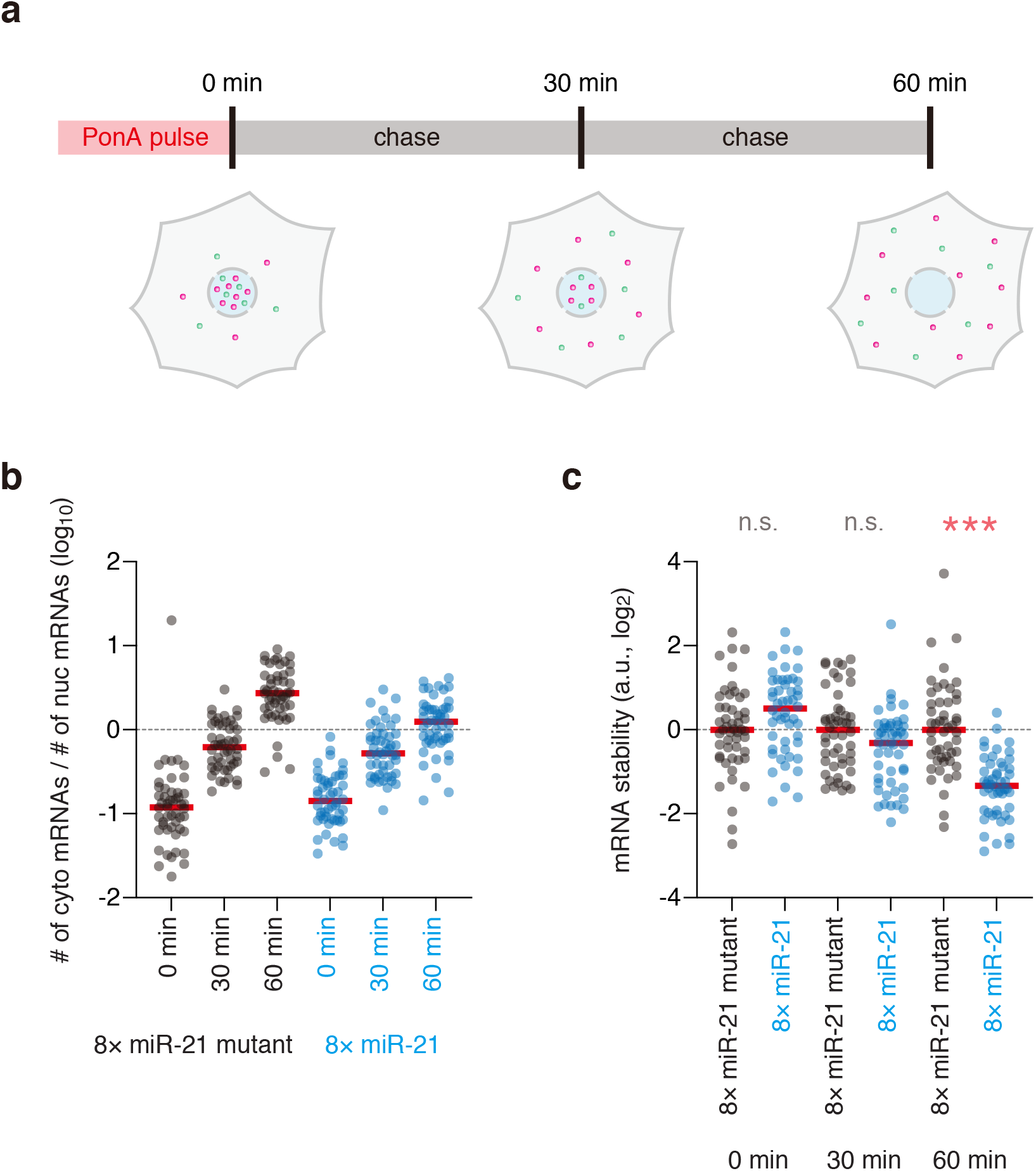
Spatiotemporal analysis of mRNA decay by single-mRNA imaging. **(a)** Schematic of spatiotemporal analysis of mRNA decay by single-mRNA imaging. In pulse-chase experiments, smFISH was performed to visualize mRNA decay at single-mRNA resolution. Green and magenta spots represent *SunTag* and *Fluc* mRNAs, respectively. **(b)** Transport of reporter mRNAs from the nucleus to the cytoplasm. In pulse-chase experiments, reporter mRNAs were labeled by smFISH. The ratios of the number of cytoplasmic mRNAs to that of nuclear mRNAs are shown. Each circle represents a single cell (n = 50 for each condition), while red lines represent the medians. cyto, cytoplasmic. nuc, nuclear. **(c)** Time-course analysis of mRNA decay by single-mRNA imaging. Images were analyzed using CellProfiler and FISH-quant. Then, mRNA stability was calculated as described in fig. S2 (see also Online Methods). Each circle represents a single cell (n = 50 for each condition), while red lines represent the medians. The results of Dunn’s multiple comparisons test are shown. *** and n.s. represent p < 0.001 and not significant (p > 0.05), respectively.

## Discussion

Historically, miRNAs have been studied primarily by ensemble methods, where a bulk collection of molecules was measured outside cells **(*8*–*10, 14*–*18, 22, 28, 29*)**. Although recent studies provided several valuable methods to analyze the function of miRNAs more precisely **(*34*–*39*)**, the behavior of individual molecules inside cells, as well as their spatiotemporal regulation, remains mostly unknown. In this study, we developed a series of single-molecule methods, which enabled us to image each step of miRNA-mediated gene silencing, i.e., RISC-binding, translational repression, and mRNA decay, inside cells. Our methods, which overcome the limitation of canonical methods, provided novel insights into when, where, and how miRNAs work inside cells (Fig. 7): 1) RISC bound to mRNAs immediately after their transport from the nucleus to the cytoplasm; 2) RISC preferred translated mRNAs over untranslated mRNAs; 3) RISC repressed translation of mRNAs within 30 min after the binding; 4) RISC reduced both the number of translated mRNAs and the number of ribosomes on translated mRNAs; 5) RISC induced mRNA decay within 60 min after the binding to mRNAs; 6) the efficiency of RISC-binding, translational repression, and mRNA decay demonstrated cell-to-cell heterogeneity. This study is complementary to a companion work by Cialek et al., where translational repression mediated by the AGO-tethering system was visualized in live cells. Their observations on translational repression are consistent with our findings 3) and 4), indicating that these results are reproducible between laboratories.

**Fig. 7.**
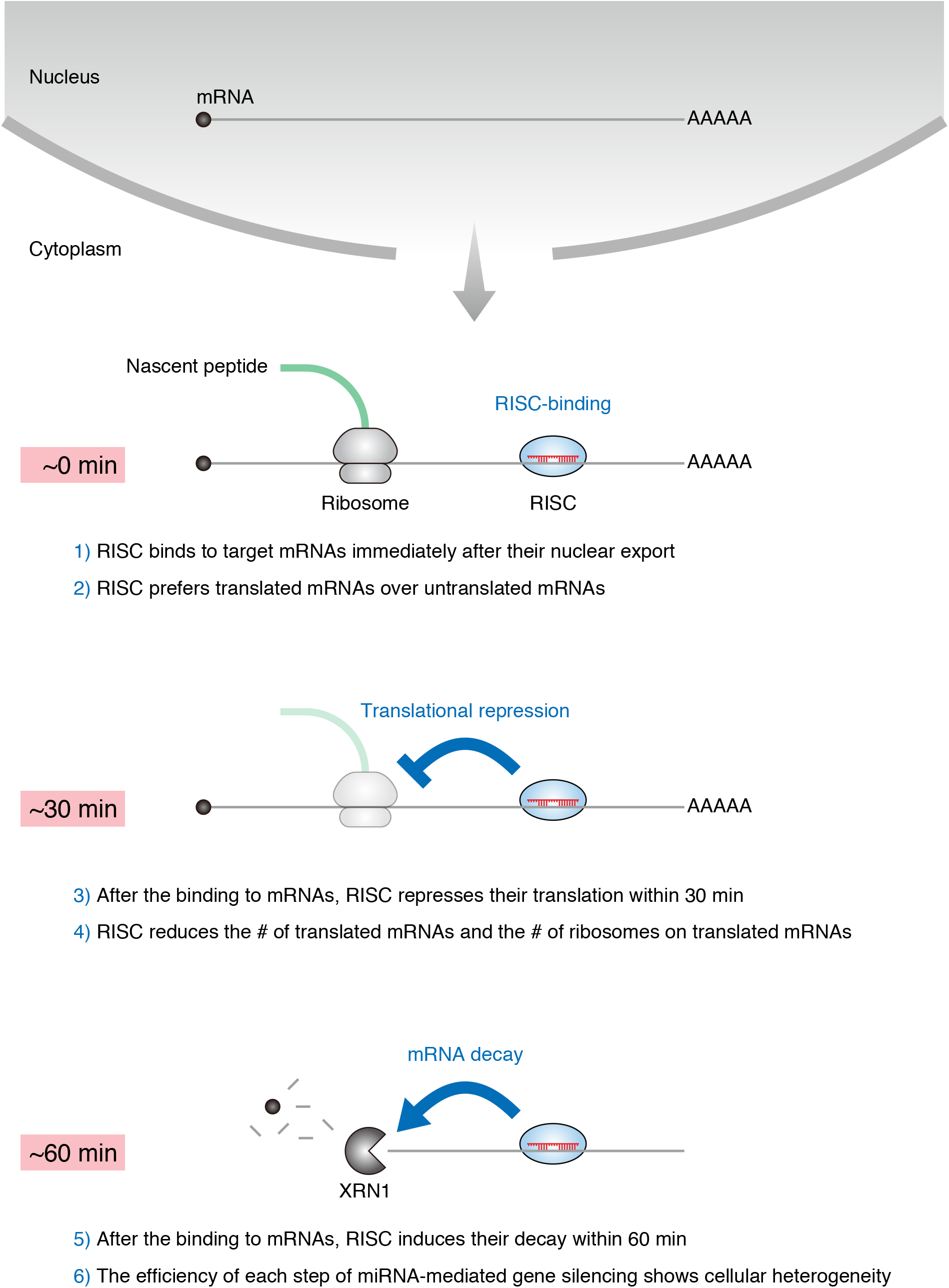
A model of miRNA-mediated gene silencing; findings from single-molecule imaging inside cells. After mRNAs are exported from the nucleus to the cytoplasm, RISC binds to them immediately. RISC preferentially binds to translated mRNAs rather than untranslated mRNAs. Then, RISC represses translation within 30 min after the binding to mRNAs. This action of RISC reduces the number of translated mRNAs inside cells, as well as the number of ribosomes on translated mRNAs. Subsequently, RISC induces mRNA decay within 60 min after the binding to mRNAs. When focusing on each cell, the efficiency of each step of miRNA-mediated gene silencing, i.e., RISC-binding, translational repression, and mRNA decay, demonstrates cell-to-cell heterogeneity. For example, miRNAs can halt the translation of target mRNAs completely within some cells, while some other cells are insensitive to the silencing.

We found that RISC preferentially bound to translated mRNAs rather than untranslated mRNAs (Fig. 4). Were RISC to recognize translated mRNAs and untranslated mRNAs with the same efficiency, this would be a waste of RISC, as RISC does not need to bind to untranslated mRNAs. Thus, we speculate that this preference would contribute to the economical use of RISC, whose number is limited inside cells (∼15,000 molecules per cell). The mechanism that enables RISC to preferentially bind to translated mRNAs should be investigated in the future.

RISC generally targets the 3′ UTR of mRNAs to recognize them. It has been proposed that this is because RISC sitting on the regions other than the 3′ UTR may be removed from mRNAs by translating ribosomes before it represses translation **(*13, 40*)**. Notably, this model is based on the assumption that RISC may need a longer time to repress translation than the speed of translation. However, the time span from RISC-binding to translational repression has been unknown. In this study, our spatiotemporal analysis revealed that RISC needs ∼30 min to repress translation after the binding to mRNAs (Fig. 5). Because this is much slower than translation initiation rates, typically faster than ∼1 per min **(*41*)**, our finding provides a missing piece to explain why RISC uses the 3′ UTR of mRNAs; RISC needs to target this ribosome-free region because translational repression by RISC is too slow to compete with ribosomes.

mRNA regulation is a fundamental step in gene regulation, thereby influencing a wide variety of biological processes and diseases. To regulate mRNAs, various RNA-binding proteins (RBPs) associate with mRNAs, which trigger the stabilization/degradation of mRNAs, the activation/repression of translation, or the translocation of mRNAs to specific areas. The methods to analyze such mRNA regulation *in situ*, however, have been limited. Therefore, our methodology, which makes it possible to visualize single mRNAs, translation, and RBP-binding simultaneously with single-molecule resolution inside cells, will be a valuable framework for studying mRNA regulation in the future.

## Online Methods

### Plasmid construction

#### The reporter plasmid for miRNA-mediated mRNA decay

The plasmid pPonA-BI-Gl NORM-LacZA TER-LacZB (Addgene plasmid # 86212), which expresses two different mRNAs (*Gl-NORM-LacZA* and *Gl-TER-LacZB*) under the control of the PonA-inducible bi-directional promoter **(*31*)**, was used as the backbone sequence. The Gl-TER-LacZB sequence was replaced by the Fluc sequence **(*42*)**, while Gl-NORM-LacZA sequence was replaced by the SunTag sequence followed by the AID degron **(*21*)**. The degron sequence was inserted in order to visualize translational repression, but not to visualize of mRNA decay. Eight miR-21 sites (or eight miR-21 mutant sites) were inserted into the 3′ UTR of the *SunTag* mRNA to recapitulate mRNA decay by miRNAs (fig. S1d). Although the backbone sequence originally had the SV40 poly(A) signals, they were replaced by the bGH poly(A) signals.

#### The reporter plasmid for miRNA-mediated translational repression

The plasmid for miRNA-mediated mRNA decay was used as the backbone sequence. The bGH poly(A) signal for the *SunTag* mRNA was replaced by the A_114_-N_40_-HhR sequence, which protects mRNAs from deadenylation and decay **(*28, 29*)**. To eliminate the unwanted RISC-binding to reporter mRNAs independent of miR-21, all potential miRNA sites (8mer, 7mer, and 6mer) of top 30 most abundant miRNAs (fig. S1, b and c) were removed from the 3′ UTR of the reporter mRNAs. Reporter plasmids were purified using NucleoBond Xtra Midi kit (MACHEREY-NAGEL, 740410) for the purpose of nucleofection.

### Cell culture

The human U2OS cells stably expressing VgEcR and RXR, which enable PonA-inducible transcription **(*32*)**, were cultured at 37°C and 5% CO_2_ in DMEM (Corning, 10-013-CV) supplemented with 10% FBS (R&D Systems, S11150H) and 1% Antibiotic-Antimycotic (Thermo Fisher, 15240-062).

### Nucleofection

U2OS cells were briefly rinsed with DPBS (Corning, 21-031-CV), followed by the treatment with Trypsin-EDTA (Thermo Fisher, 25300-054) to detach them. After adding culture media to neutralize trypsinization, cells were centrifuged at 300 g for 1 min. The pellets (approx. 1 × 10^6^ cells) were resuspended with 100 μl of Ingenio Electroporation Solution (Mirus, MIR50115) containing 2 μg of reporter plasmids. Then, nucleofection was performed in the electroporation cuvette (Mirus, MIR50115) using Nucleofector II (Lonza). Nucleofected cells were cultured on the coverslips (Thermo Fisher, 12-545-81) coated with collagen (Cell Applications, 125-50) in culture media.

### Drug treatment

One day after nucleofection, U2OS cells were treated with 20 μM PonA (Santa Cruz, sc-202768A) for 30 min to induce transcription of reporter mRNAs. After washing cells with culture media, cells were incubated for 1 hr. Subsequently, cells were fixed with 4% paraformaldehyde (Electron Microscopy Sciences, 15714) in 1× PBS (MilliporeSigma, 11666789001) for 10 min. For puromycin experiments (fig. S5), cells were treated with 100 μg/ml puromycin (MilliporeSigma, CAS 58-58-2) from the beginning of PonA treatment until fixation.

### Pulse-chase experiment

One day after nucleofection, U2OS cells were treated with 20 μM PonA for 30 min to induce transcription of reporter mRNAs. After washing cells with culture media, cells were fixed at 0, 30, and 60 min after PonA treatment with 4% paraformaldehyde in 1× PBS for 10 min.

### smFISH

Fixed cells were permeabilized with 0.1% Triton X-100 (MilliporeSigma, T9284) in 1× PBS for 10 min, followed by washing with 1× PBS. Subsequently, cells were incubated with the pre-hybridization solution containing 10% deionized formamide (Thermo Fisher, AC205821000), 2× SSC (MilliporeSigma, 11666681001), 0.5% UltraPure BSA (Thermo Fisher, AM2618), and 40 U/ml SUPERase In RNase Inhibitor (Thermo Fisher, AM2696) for 30 min. Then, cells were incubated with the hybridization solution containing 10% deionized formamide, 2× SSC, 10% dextran sulfate (MilliporeSigma, D8906), 1 mg/ml competitor tRNA (MilliporeSigma, 10109541001), 0.05% UltraPure BSA, 40 U/ml SUPERase In RNase Inhibitor, and 50 nM smFISH probes for 3 hr at 37°C. After washing with 10% deionized formamide in 2× SSC, followed by washing with 2× SSC, coverslips were mounted onto glass slides (Thermo Fisher, 3051-002) using ProLong Diamond Antifade Mountant with DAPI (Thermo Fisher, P36962). The smFISH probes toward *Fluc* mRNAs were designed using Stellaris Probe Designer version 4.2 (Biosearch Technologies). The smFISH probes conjugated with Quasar 570 toward *Fluc* mRNAs and the smFISH probes conjugated with Quasar 670 toward *SunTag* mRNAs **(*21*)** were synthesized by Biosearch Technologies. The sequences of smFISH probes are listed in the Table S1.

### SINAPS

Fixed cells were permeabilized and pre-hybridized as described in the smFISH section. Then, cells were incubated with the hybridization solution containing 10% deionized formamide, 2× SSC, 10% dextran sulfate, 1 mg/ml competitor tRNA, 0.05% UltraPure BSA, 40 U/ml SUPERase In RNase Inhibitor, 50 nM smFISH probes conjugated with Quasar 570 toward *SunTag* mRNAs **(*21*)**, and 10 μg/ml anti-GCN4 Rabbit antibody (Absolute Antibody, AB00436-23.0) for 3 hr at 37°C. Subsequently, cells were washed with 10% deionized formamide in 2× SSC, followed by incubation with 10% deionized formamide in 2× SSC supplemented with 2 μg/ml Goat anti-Rabbit IgG conjugated with Alexa Fluor 488 (Thermo Fisher, A-11034) for 30 min at 37°C. After washing with 2× SSC, coverslips were mounted onto glass slides using ProLong Diamond Antifade Mountant with DAPI.

### IF-FISH

Fixed cells were permeabilized with 0.1% Triton X-100 in 1× PBS, followed by washing with 1× PBS. Subsequently, cells were incubated with the blocking buffer (1× PBS, 0.02% Triton X-100, 0.5% UltraPure BSA, and 40 U/ml SUPERase In RNase Inhibitor) for 30 min. Then, cells were incubated with the blocking buffer supplemented with 24 μg/ml anti-AGO2 Mouse antibody (FUJIFILM Wako Pure Chemical, 015-22031) for 1 hr. Cells were washed with the blocking buffer, followed by incubation with the blocking buffer supplemented with 2 μg/ml Goat anti-Mouse IgG conjugated with Alexa Fluor 647 (Thermo Fisher, A-21236) for 30 min. After washing with 1× PBS, immunostained cells were pre-hybridized and hybridized to perform smFISH as described in the smFISH section. For the experiments to visualize single mRNAs, translation, and RISC-binding simultaneously (Figs. 4 and 5), immunostained cells were pre-hybridized and hybridized to perform SINAPS as described in the SINAPS section.

### Image acquisition

Slides were imaged on the BX63 automated wide-field fluorescence microscope (Olympus) equipped with the SOLA FISH light engines (Lumencor), the ORCA-R2 cooled digital CCD camera (Hamamatsu Photonics), the 60× 1.35 NA super apochromat objective (Olympus, UPLSAPO60XO), and zero pixel shift filter sets: DAPI-5060C-Zero, FITC-5050A-Zero, Cy3-4040C-Zero, and Cy5-4040C-Zero (Semrock). To acquire multi-color 3D images, the microscope was controlled with MetaMorph software (Molecular Devices), where the Multi Dimensional Acquisition mode was selected. Exposure times for each color were 100 ms for CY5 (gain: 2), 100 ms for CY3 (gain: 2), 100 ms for FITC (gain: 2), and 5 ms for DAPI (gain: 0). For each color, Z stacks spanning the entire volume of cells were acquired by imaging every 200 nm along the z-axis. Image pixel size: XY, 107.5 nm; Z, 200 nm. n = 50 cells for each experiment.

### Image analysis

Images were analyzed using FISH-quant **(*25*)**, an algorithm implemented in MATLAB. Briefly, after background subtraction, FISH-quant automatically detects fluorescent spots and localizes them in 3D at sub-pixel resolution by fitting 3D Gaussians. This provides the number of spots inside cells, the intensity of each spot, and the 3D position of each spot **(*25*)**. To distinguish nuclear and cytoplasmic areas, the nuclei and cells were visualized by DAPI and the non-specific background signals of smFISH probes, respectively. Their outlines were automatically detected by the CellProfiler algorithm **(*33*)**, followed by conversion to the outline files compatible with FISH-quant. Based on these outlines, all spots were classified into “nuclear” and “cytoplasmic”. To prepare images for figures (Figs. 1c, 2c, 3b, and 4b), raw images were processed using ImageJ software (Version: 2.1.0/1.53c).

### Colocalization analysis

Colocalization in 3D was analyzed using FISH-quant **(*25*)**. First, the average drift between different colors was calculated and corrected. Then, using the 3D positions of spots, their 3D distances were calculated. When two spots in different colors were localized within the maximum allowed distance (mRNA-SunTag, 500 nm; mRNA-AGO, 250 nm), these spots were considered colocalized. Based on the 3D colocalization with SunTag and AGO, mRNAs were classified into “translated”, “untranslated”, “RISC-bound”, and “RISC-unbound”. Likewise, SunTag and AGO spots were also classified into “on mRNAs” and “free”, depending on the colocalization with mRNAs.

### Data analysis

#### Profiling miRNAs expressed in U2OS cells

The small RNA-seq data of U2OS cells (GEO accession: GSM416754) **(*22*)** was reanalyzed as described previously **(*42*)**.

#### Profiling AGO proteins expressed in U2OS cells

From the proteome data of U2OS cells **(*30*)**, the copies per cell values of AGO1 (UniProtKB: Q9UL18), AGO2 (UniProtKB: Q9UKV8), AGO3 (UniProtKB: Q9H9G7), and AGO4 (UniProtKB: Q9HCK5) were extracted.

#### Data analysis for miRNA-mediated mRNA decay

The mRNA stability of each cell, which is used for single cell analysis (Figs. 1e and 6c), was calculated by Equation 1, where *S(k)* is the mRNA stability of the *k*^*th*^ cell. *M*_*Fluc,cyto*_*(k)* and *M*_*Sun,cyto*_*(k)* are the number of *Fluc* mRNAs and *SunTag* mRNAs, respectively, in the cytoplasm of the *k*^*th*^ cell.

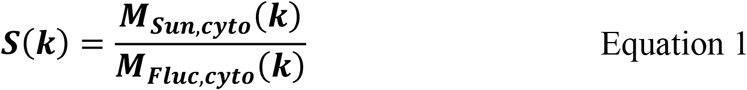

The mRNA stability of the cell population (50 cells), *S*_*bulk*_, which is used for bulk analysis (Fig. 1d), was calculated by Equation 2.

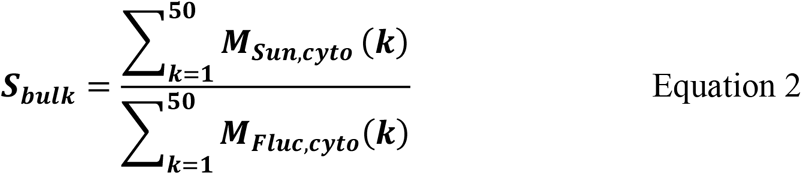

#### Data analysis for miRNA-mediated translational repression

The translational efficiency of each cell, which is used for single cell analysis (Figs. 2e and 5e), was calculated by Equation 3, where *T*_*eff*_*(k)* is the translational efficiency of the *k*^*th*^ cell. *I*_*Sun,coloc,cyto*_*(k)* is the total intensity of SunTag spots on mRNAs in the cytoplasm of the *k*^*th*^ cell.

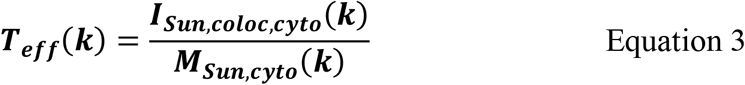

The translational efficiency of the cell population (50 cells), *T*_*eff,bulk*_, which is used for bulk analysis (Fig. 2d), was calculated by Equation 4.

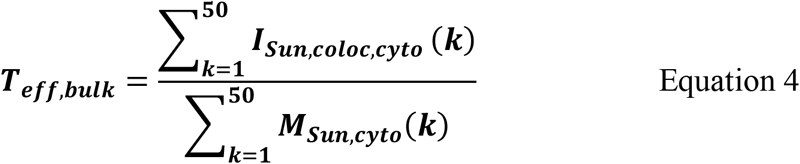

The fraction of translated mRNAs of each cell, which is used for single cell analysis (Fig. 2f and S10b), was calculated by Equation 5, where *T*_*fra*_*(k)* is the fraction of translated mRNAs of the *k*^*th*^ cell. *M*_*Sun,coloc,cyto*_*(k)* is the number of translated *SunTag* mRNAs in the cytoplasm of the *k*^*th*^ cell.

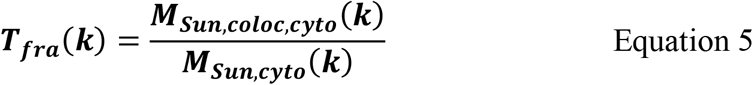

The number of ribosomes on translated mRNAs, *R*, which is used for single-molecule analysis (Figs. 2i and 5f), was calculated by Equation 6. *i*_*MED,Sun,free,cyto*_ is the median of the intensities of free SunTag spots in the cytoplasm, while *i*_*Sun,coloc,cyto*_ is the intensity of each SunTag spot on mRNAs in the cytoplasm. Bright SunTag spots on mRNAs should contain partial SunTag peptides, which had not been fully translated, as well as full-length SunTag peptides. Thus, the exact number of ribosomes on translated mRNAs should be larger than our values **(*21*)**.

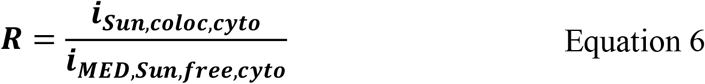

#### Data analysis for RISC-binding

The RISC-binding efficiency of each cell, which is used for single cell analysis (Figs. 3d and 5c), was calculated by Equation 7, where *A*_*eff*_*(k)* is the RISC-binding efficiency of the *k*^*th*^ cell. *I*_*AGO,coloc,cyto*_*(k)* is the total intensity of AGO spots on mRNAs in the cytoplasm of the *k*^*th*^ cell.

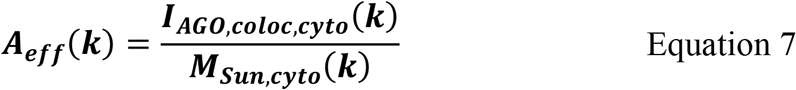

The RISC-binding efficiency of the cell population (50 cells), *A*_*eff,bulk*_, which is used for bulk analysis (Fig. 3c), was calculated by Equation 8.

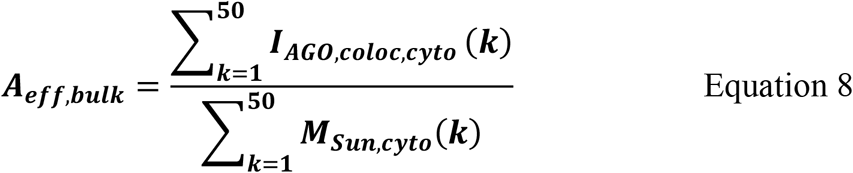

The fraction of RISC-bound mRNAs of each cell, which is used for single cell analysis (Figs. 3e and S10a), was calculated by Equation 9, where *A*_*fra*_*(k)* is the fraction of RISC-bound mRNAs of the *k*^*th*^ cell. *M*_*AGO,coloc,cyto*_*(k)* is the number of RISC-bound *SunTag* mRNAs in the cytoplasm of the *k*^*th*^ cell.

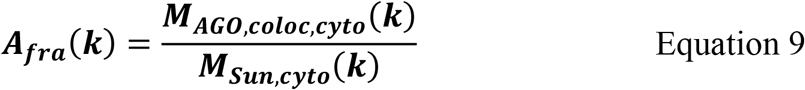

#### Data analysis for mRNA export

The mRNA export efficiency of each cell, which is used for single cell analysis (Figs. 5b and 6b), was calculated by Equation 10, where *E(k)* is the mRNA export efficiency of the *k*^*th*^ cell. *M*_*Sun,nuc*_*(k)* and *M*_*Sun,cyto*_*(k)* are the number of *SunTag* mRNAs in the nucleus and the cytoplasm, respectively, of the *k*^*th*^ cell.

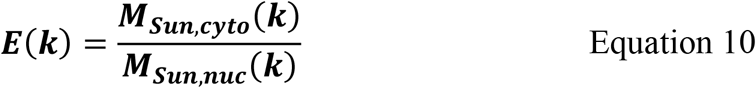

### Statistical analysis

Mann Whitney tests were performed in Figs. 1e, 2e, 2f, 2i, 3d, 3e, S3a, S3b, S5b, S5c, S6a, S6b, and S8b, while Dunn’s multiple comparisons tests were performed in Figs. 2h, 3g, 5c, 5d, 5e, 5f, 6c, S10a, and S10b. *** and n.s. represent p < 0.001 and not significant (p > 0.05), respectively, in Figs. 5c, 5e, 6c, S10a, and S10b. These statistical tests, calculation of Pearson correlation coefficient (r) (Figs. 4d and S3c), and simple linear regression (fig. S3c) were performed using GraphPad Prism (Version: 8), which is also used to create all graphs in this study.

### Data availability statement

All data generated and analyzed during this study are available from the corresponding authors on reasonable request.

## Acknowledgements

We are grateful to Yukihide Tomari for providing pAWH-Rluc-let-7-A_114_-N_40_-HhR. We thank Xiuhua Meng and Melissa Lopez-Jones for technical assistance, Hanae Sato for sharing the protocols of smFISH and SINAPS, and Carolina Eliscovich for sharing the protocol for IF-FISH. We are grateful to Keisuke Shoji for supporting the analysis of small RNA-seq data. We thank all the members of the Robert H. Singer laboratory for insightful discussion and critical comments. The authors thank Cialek et al. for sharing their preliminary data. This work was supported by NIH grants R01NS083085 and R35GM136296 (to R.H.S.), JSPS Overseas Research Fellowships (to H.K.), JBS Osamu Hayaishi Memorial Scholarship for Study Abroad (to H.K.), and JST PRESTO Grant JPMJPR20E7 (to H.K.).

## Author Contributions

H.K. and R.H.S. designed the project. H.K. performed experiments and analyzed data. H.K. and R.H.S. wrote the manuscript.

