## Supplementary Information for "Single-molecule imaging of microRNA-mediated gene silencing in cells"

##### Supplementary Figure Legends

**Fig. S1. Expression profile of miRNAs in U2OS cells.**

(**a** to **c**) The expression profile of miRNAs in U2OS cells. The small RNA-seq data (Mayr and Bartel, 2009) was reanalyzed. The length of small RNAs (a), the number of reads of each miRNA (b), and their relative occupancies (c) are shown. Top 30 most abundant miRNAs, whose sites were removed from reporter mRNAs, occupy ~80% of total population.

(**d**) The sequences of the miR-21 site (left) and the miR-21 mutant site (right). The miR-21 site was designed so that the positions 2–8 (seed region) and 13–16 (3′ supplemental region) of miR-21 form base-pairs with the miR-21 site. The miR-21 mutant site does not form base-pairs with miR-21.

**Fig. S2. Workflow for single-molecule imaging of miRNA-mediated mRNA decay.**

The workflow for single-molecule imaging of miRNA-mediated mRNA decay is shown step by step. Detailed methods are provided in Online Methods.

**Fig. S3. Supplemental data for Fig. 1.**

(**a** and **b**) The number of *Fluc* mRNAs (a) and *SunTag* mRNAs (b) detected in U2OS cells. Images were analyzed using CellProfiler and FISH-quant. Each circle represents a single cell (n = 50 for each condition), while red lines represent the medians. The p values of Mann Whitney test are shown. n.s., not significant.

(**c**) Positive correlation between the number of *Fluc* mRNAs and *SunTag* mRNAs. Each circle represents a single cell (n = 50), while the red line represents the result of simple linear regression. The Pearson correlation coefficient (r) is shown.

**Fig. S4. Workflow for single-molecule imaging of miRNA-mediated translational repression.**

The workflow for single-molecule imaging of miRNA-mediated translational repression is shown step by step. Detailed methods are provided in Online Methods.

**Fig. S5. Validation of SINAPS experiments by puromycin treatment.**

(**a** and **b**) Reduction of translational efficiency by puromycin treatment. Images were analyzed using CellProfiler and FISH-quant. Then, translational efficiency was calculated as described in fig. S4 (see also Online Methods). The results of bulk analysis (a) and single-cell analysis (b) are shown. In (b), each circle represents a single cell (n = 50 for each condition), while red lines represent the medians. The p value of Mann Whitney test is shown. The puromycin – data in (a) and (b) are identical to the data in Fig. 2d and 2e, respectively.

(**c**) Reduction of the fraction of translated mRNAs by puromycin treatment. The fraction of translated mRNAs was calculated as described in fig. S4 (see also Online Methods). Each circle represents a single cell (n = 50 for each condition), while red lines represent the medians. The p value of Mann Whitney test is shown. The puromycin – data are identical to the data in Fig. 2f.

**Fig. S6. Supplemental data for Fig. 2.**

(**a** and **b**) The number of reporter mRNAs (a) and SunTag spots (b) detected in U2OS cells. Images were analyzed using CellProfiler and FISH-quant. Each circle represents a single cell (n = 50 for each condition), while red lines represent the medians. The p values of Mann Whitney test are shown. n.s., not significant.

**Fig. S7. Workflow for single-molecule imaging of RISC-binding.**

The workflow for single-molecule imaging of RISC-binding is shown step by step. Detailed methods are provided in Online Methods.

**Fig. S8. Supplemental data for Fig. 3.**

(**a**) The expression profile of AGO proteins in U2OS cells. The proteome data of U2OS cells (Beck et al., 2011) was reanalyzed. The number of copies per cell are shown. N.D., not detected. In U2OS cells, AGO2 is predominantly expressed.

(**b**) The number of AGO spots detected in U2OS cells. Images were analyzed using CellProfiler and FISH-quant. Each circle represents a single cell (n = 50 for each condition), while red lines represent the medians. The p value of Mann Whitney test is shown. n.s., not significant.

**Fig. S9. Workflow for simultaneous visualization of single mRNAs, translation, and RISC-binding.**

The workflow for simultaneous visualization of single mRNAs, translation, and RISC-binding is shown step by step. Detailed methods are provided in Online Methods.

**Fig. S10. Supplemental data for Fig. 5.**

(**a** and **b**) Time-course analysis of RISC-binding (a) and translational repression (b) by single-mRNA imaging. Images were analyzed using CellProfiler and FISH-quant. Then, the fraction of RISC-bound mRNAs (a) and of translated mRNAs (b) were calculated as described in fig. S9 (see also Online Methods). Each circle represents a single cell (n = 50 for each condition), while red lines represent the medians. The results of Dunn's multiple comparisons test are shown. *** and n.s. represent $p < 0.001$ and not significant ($p > 0.05$), respectively.

(**c**) The ratio of RISC-unbound untranslated (magenta), RISC-unbound translated (green), RISC-bound untranslated (cyan), and RISC-bound translated (orange) mRNAs. All mRNAs were classified into these four classes based on 3D-colocalization analysis.

#### Supplementary Table Legends

**Table S1. The sequences of smFISH probes used in this study.**

The sequences of smFISH probes toward *Fluc* mRNAs and *SunTag* mRNAs used in this study are listed.

Kobayashi & Singer, Fig. S1

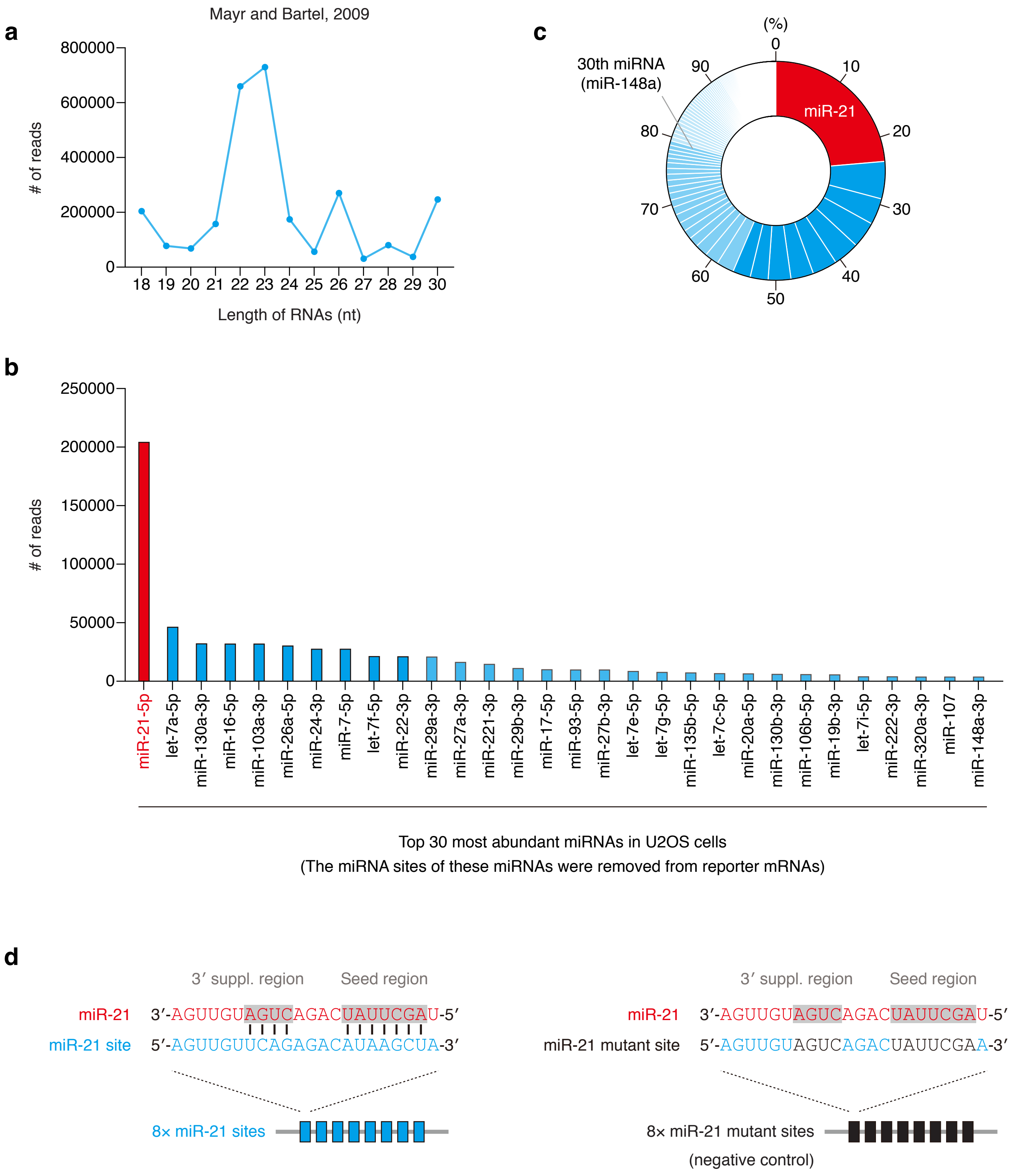

**Fig. S1. Expression profile of miRNAs in U2OS cells.**

(**a** to **c**) The expression profile of miRNAs in U2OS cells. The small RNA-seq data (Mayr and Bartel, 2009) was reanalyzed. The length of small RNAs (a), the number of reads of each miRNA (b), and their relative occupancies (c) are shown. Top 30 most abundant miRNAs, whose sites were removed from reporter mRNAs, occupy ~80% of total population.

(**d**) The sequences of the miR-21 site (left) and the miR-21 mutant site (right). The miR-21 site was designed so that the positions 2–8 (seed region) and 13–16 (3′ supplemental region) of miR-21 form base-pairs with the miR-21 site. The miR-21 mutant site does not form base-pairs with miR-21.

**1. Induction of transcription**

**2. smFISH**

Label *Fluc* mRNAs by smFISH probes (Quasar 570)

Label *SunTag* mRNAs by smFISH probes (Quasar 670)

Label nuclei by DAPI

**3. Image acquisition**

3-color: *Fluc* mRNAs (orange), *SunTag* mRNAs (far red), nuclei (blue)

3D: pixel size: XY, 107.5 nm; Z, 200 nm

**4. Image analysis**

Detect the outlines of cells and nuclei (by CellProfiler)

Detect spots of *Fluc* mRNAs and *SunTag* mRNAs (by FISH-quant)

Localize spots in 3D at sub-picxel resolution by fitting 3D Gaussians

Extract data of cytoplasmic spots: #, intensities, positions in X, Y, Z

**5. Data analysis**

$$\text{mRNA stability} = \frac{\#\text{ of } \textit{SunTag} \text{ mRNAs}}{\#\text{ of } \textit{Fluc} \text{ mRNAs}}$$

**Fig. S2. Workflow for single-molecule imaging of miRNA-mediated mRNA decay.**

The workflow for single-molecule imaging of miRNA-mediated mRNA decay is shown step by step. Detailed methods are provided in Online Methods.

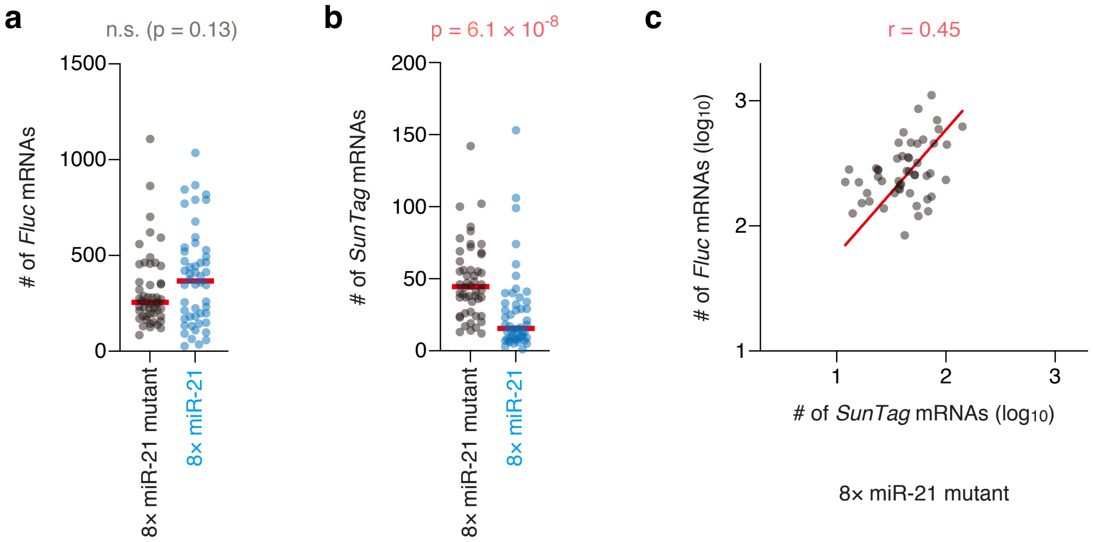
a
n.s. (p = 0.13)
### of Fluc mRNAs
1500
1000
500
0
8× miR-21 mutant
8× miR-21
b
p = 6.1 × 10-8
### of SunTag mRNAs
200
150
100
50
0
8× miR-21 mutant
8× miR-21
c
r = 0.45
### of Fluc mRNAs (log10)
3
2
1
1
2
3
### of SunTag mRNAs (log10)
8× miR-21 mutant

**Fig. S3. Supplemental data for Fig. 1.**

(**a** and **b**) The number of *Fluc* mRNAs (a) and *SunTag* mRNAs (b) detected in U2OS cells. Images were analyzed using CellProfiler and FISH-quant. Each circle represents a single cell (n = 50 for each condition), while red lines represent the medians. The p values of Mann Whitney test are shown. n.s., not significant. (**c**) Positive correlation between the number of *Fluc* mRNAs and *SunTag* mRNAs. Each circle represents a single cell (n = 50), while the red line represents the result of simple linear regression. The Pearson correlation coefficient (r) is shown.

Kobayashi & Singer, Fig. S4

**1. Induction of transcription**

**2. SINAPS**

Label SunTag peptides by anti-GCN4 antibodies (Alexa 488)

Label *SunTag* mRNAs by smFISH probes (Quasar 570)

Label nuclei by DAPI

**3. Image acquisition**

3-color: SunTag (green), mRNAs (orange), nuclei (blue)

3D: pixel size: XY, 107.5 nm; Z, 200 nm

**4. Image analysis**

Detect the outlines of cells and nuclei (by CellProfiler)

Detect spots of SunTag and mRNAs (by FISH-quant)

Localize spots in 3D at sub-picxel resolution by fitting 3D Gaussians

Extract data of cytoplasmic spots: #, intensities, positions in X, Y, Z

**5. Colocalization analysis**

Analyse colocalization based on the 3D distance between spots

Classify mRNAs into “untranslated” and “translated”

Classify SunTag into “free” and “on mRNAs”

**6. Data analysis**

$$\text{Translational efficiency} = \frac{\text{Intensity of SunTag on mRNAs}}{\text{\# of mRNAs}}$$

$$\text{Fraction of translated mRNAs} = \frac{\text{\# of translated mRNAs}}{\text{\# of mRNAs}}$$

$$\text{\# of ribosomes} = \frac{\text{Intensity of SunTag on mRNAs}}{\text{Intensity of free SunTag}}$$

**Fig. S4. Workflow for single-molecule imaging of miRNA-mediated translational repression.**

The workflow for single-molecule imaging of miRNA-mediated translational repression is shown step by step. Detailed methods are provided in Online Methods.

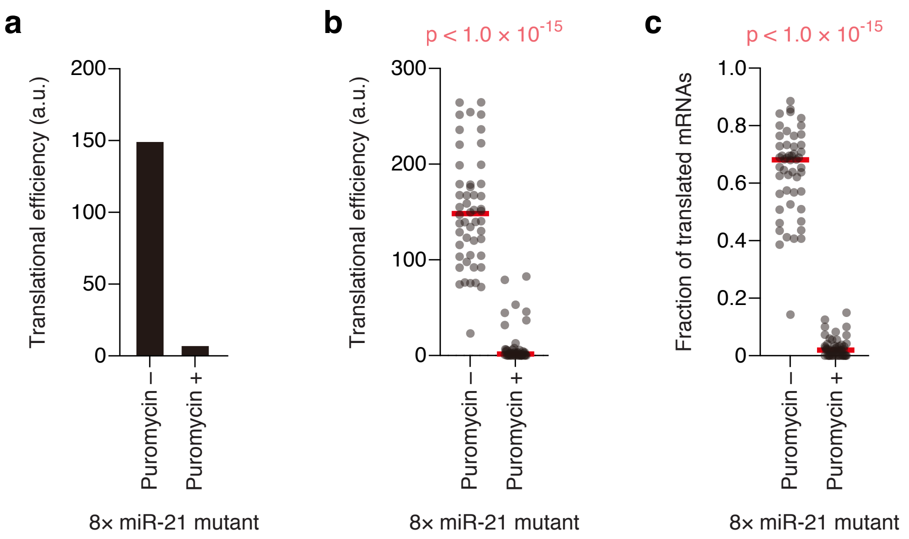
a
b
c
Translational efficiency (a.u.)
200
150
100
50
0
Puromycin –
Puromycin +
8× miR-21 mutant
p < 1.0 × 10-15
Translational efficiency (a.u.)
300
200
100
0
Puromycin –
Puromycin +
8× miR-21 mutant
p < 1.0 × 10-15
Fraction of translated mRNAs
1.0
0.8
0.6
0.4
0.2
0
Puromycin –
Puromycin +
8× miR-21 mutant

**Fig. S5. Validation of SINAPS experiments by puromycin treatment.**

(**a** and **b**) Reduction of translational efficiency by puromycin treatment. Images were analyzed using CellProfiler and FISH-quant. Then, translational efficiency was calculated as described in fig. S4 (see also Online Methods). The results of bulk analysis (a) and single-cell analysis (b) are shown. In (b), each circle represents a single cell (n = 50 for each condition), while red lines represent the medians. The p value of Mann Whitney test is shown. The puromycin – data in (a) and (b) are identical to the data in Fig. 2d and 2e, respectively.

(**c**) Reduction of the fraction of translated mRNAs by puromycin treatment. The fraction of translated mRNAs was calculated as described in fig. S4 (see also Online Methods). Each circle represents a single cell (n = 50 for each condition), while red lines represent the medians. The p value of Mann Whitney test is shown. The puromycin – data are identical to the data in Fig. 2f.

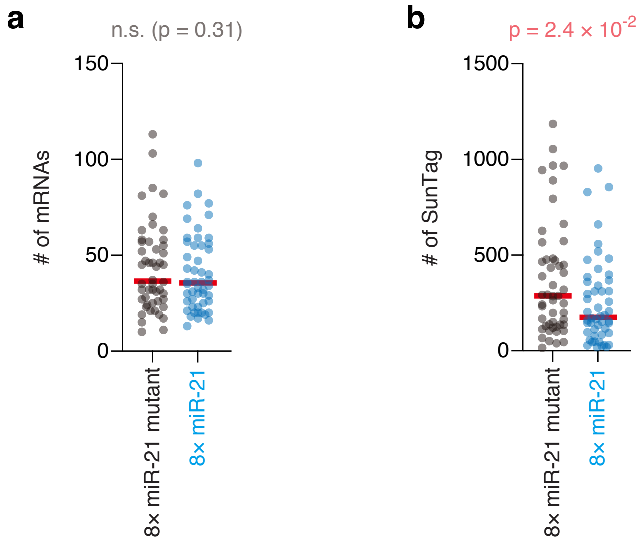
a
n.s. (p = 0.31)
150
100
50
0
### of mRNAs
8× miR-21 mutant
8× miR-21
b
p = 2.4 × 10-2
1500
1000
500
0
### of SunTag
8× miR-21 mutant
8× miR-21

**Fig. S6. Supplemental data for Fig. 2.**

(**a** and **b**) The number of reporter mRNAs (a) and SunTag spots (b) detected in U2OS cells. Images were analyzed using CellProfiler and FISH-quant. Each circle represents a single cell (n = 50 for each condition), while red lines represent the medians. The p values of Mann Whitney test are shown. n.s., not significant.

**1. Induction of transcription**

**2. IF-FISH**

Label *SunTag* mRNAs by smFISH probes (Quasar 570)

Label RISC by anti-AGO antibodies (Alexa 647)

Label nuclei by DAPI

**3. Image acquisition**

3-color: mRNAs (orange), RISC (far red), nuclei (blue)

3D: pixel size: XY, 107.5 nm; Z, 200 nm

**4. Image analysis**

Detect the outlines of cells and nuclei (by CellProfiler)

Detect spots of mRNAs and RISC (by FISH-quant)

Localize spots in 3D at sub-picxel resolution by fitting 3D Gaussians

Extract data of cytoplasmic spots: #, intensities, positions in X, Y, Z

**5. Colocalization analysis**

Analyse colocalization based on the 3D distance between spots

Classify mRNAs into “RISC-unbound” and “RISC-bound”

Classify RISC into “free” and “on mRNAs”

**6. Data analysis**

$$\text{RISC-binding efficiency} = \frac{\text{Intensity of RISC on mRNAs}}{\text{\# of mRNAs}}$$

$$\text{Fraction of RISC-bound mRNAs} = \frac{\text{\# of RISC-bound mRNAs}}{\text{\# of mRNAs}}$$

**Fig. S7. Workflow for single-molecule imaging of RISC-binding.**

The workflow for single-molecule imaging of RISC-binding is shown step by step. Detailed methods are provided in Online Methods.

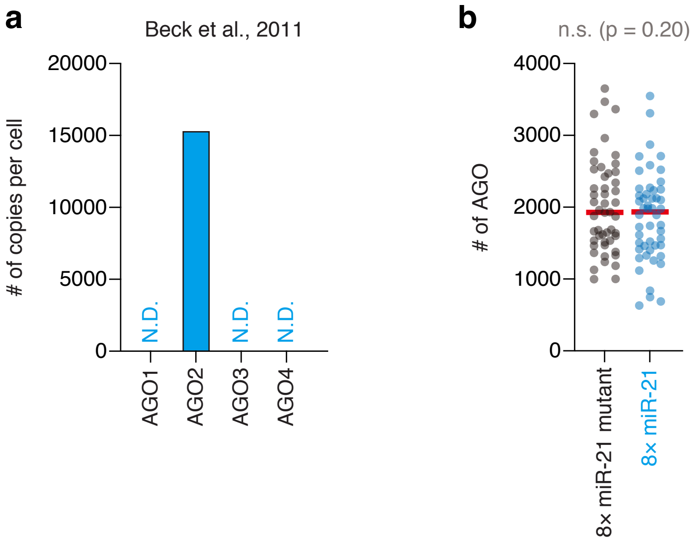
a
Beck et al., 2011
### of copies per cell
20000
15000
10000
5000
0
N.D.
N.D.
N.D.
AGO1
AGO2
AGO3
AGO4
b
n.s. (p = 0.20)
### of AGO
4000
3000
2000
1000
0
8× miR-21 mutant
8× miR-21

**Fig. S8. Supplemental data for Fig. 3.**

(**a**) The expression profile of AGO proteins in U2OS cells. The proteome data of U2OS cells (Beck et al., 2011) was reanalyzed. The number of copies per cell are shown. N.D., not detected. In U2OS cells, AGO2 is predominantly expressed.

(**b**) The number of AGO spots detected in U2OS cells. Images were analyzed using CellProfiler and FISH-quant. Each circle represents a single cell (n = 50 for each condition), while red lines represent the medians. The p value of Mann Whitney test is shown. n.s., not significant.

**1. Induction of transcription**

**2. SINAPS + IF-FISH**

Label SunTag peptides by anti-GCN4 antibodies (Alexa 488)

Label *SunTag* mRNAs by smFISH probes (Quasar 570)

Label RISC by anti-AGO antibodies (Alexa 647)

Label nuclei by DAPI

**3. Image acquisition**

4-color: SunTag (green), mRNAs (orange), RISC (far red), nuclei (blue)

3D: pixel size: XY, 107.5 nm; Z, 200 nm

**4. Image analysis**

Detect the outlines of cells and nuclei (by CellProfiler)

Detect spots of SunTag, mRNAs, and RISC (by FISH-quant)

Localize spots in 3D at sub-picxel resolution by fitting 3D Gaussians

Extract data of cytoplasmic spots: #, intensities, positions in X, Y, Z

**5. Colocalization analysis**

Analyse colocalization based on the 3D distance between spots

Classify mRNAs into “untranslated” and “translated”

Classify SunTag into “free” and “on mRNAs”

Classify mRNAs into “RISC-unbound” and “RISC-bound”

Classify RISC into “free” and “on mRNAs”

**6. Data analysis**

$$\text{Translational efficiency} = \frac{\text{Intensity of SunTag on mRNAs}}{\text{\# of mRNAs}}$$

$$\text{Fraction of translated mRNAs} = \frac{\text{\# of translated mRNAs}}{\text{\# of mRNAs}}$$

$$\text{\# of ribosomes} = \frac{\text{Intensity of SunTag on mRNAs}}{\text{Intensity of free SunTag}}$$

$$\text{RISC-binding efficiency} = \frac{\text{Intensity of RISC on mRNAs}}{\text{\# of mRNAs}}$$

$$\text{Fraction of RISC-bound mRNAs} = \frac{\text{\# of RISC-bound mRNAs}}{\text{\# of mRNAs}}$$

**Fig. S9. Workflow for simultaneous visualization of single mRNAs, translation, and RISC-binding.** The workflow for simultaneous visualization of single mRNAs, translation, and RISC-binding is shown step by step. Detailed methods are provided in Online Methods.

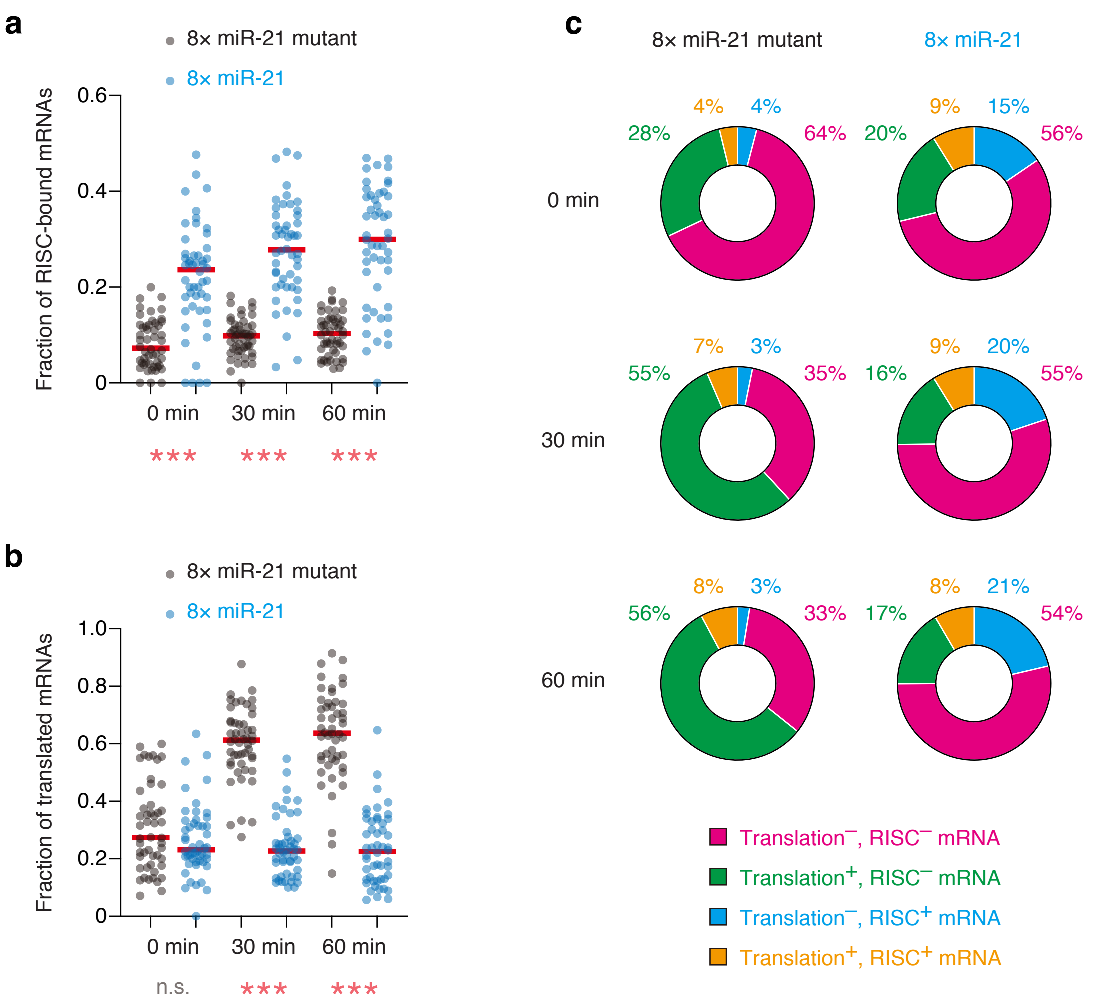
a
8× miR-21 mutant
8× miR-21
Fraction of RISC-bound mRNAs
0.6
0.4
0.2
0
0 min
30 min
60 min
***
***
***
b
8× miR-21 mutant
8× miR-21
Fraction of translated mRNAs
1.0
0.8
0.6
0.4
0.2
0
0 min
30 min
60 min
n.s.
***
***
c
8× miR-21 mutant
8× miR-21
0 min
28%
4%
4%
64%
20%
9%
15%
56%
30 min
55%
7%
3%
35%
16%
9%
20%
55%
60 min
56%
8%
3%
33%
17%
8%
21%
54%
Translation⁻, RISC⁻ mRNA
Translation⁺, RISC⁻ mRNA
Translation⁻, RISC⁺ mRNA
Translation⁺, RISC⁺ mRNA

**Fig. S10. Supplemental data for Fig. 5.**

(**a** and **b**) Time-course analysis of RISC-binding (a) and translational repression (b) by single-mRNA imaging. Images were analyzed using CellProfiler and FISH-quant. Then, the fraction of RISC-bound mRNAs (a) and of translated mRNAs (b) were calculated as described in fig. S9 (see also Online Methods). Each circle represents a single cell ($n = 50$ for each condition), while red lines represent the medians. The results of Dunn's multiple comparisons test are shown. *** and n.s. represent $p < 0.001$ and not significant ($p > 0.05$), respectively.

(**c**) The ratio of RISC-unbound untranslated (magenta), RISC-unbound translated (green), RISC-bound untranslated (cyan), and RISC-bound translated (orange) mRNAs. All mRNAs were classified into these four classes based on 3D-colocalization analysis.

**Table S1. The sequences of smFISH probes used in this study.**

smFISH probes toward *Fluc*

| Sequence Name | Sequence |
|---|---|
| Fluc_1 | catcggtgaaggcaatggtg |
| Fluc_2 | taggtgatgtccacctcaat |
| Fluc_3 | cgcacagacatctcgaagta |
| Fluc_4 | tctcagagcacaccacgatg |
| Fluc_5 | cactggcatgaagaactgca |
| Fluc_6 | cactccgatgaacagggcac |
| Fluc_7 | gtaaatgtcgttagcagggg |
| Fluc_8 | cttagacacgaacaccacgg |
| Fluc_9 | gtccatgatgatgatcttct |
| Fluc_10 | cgaatgtgtacatgctctgg |
| Fluc_11 | ctggcacgaagtcgtactcg |
| Fluc_12 | gttttgtccctgtcgaaaga |
| Fluc_13 | cagagctgttcatgatcagg |
| Fluc_14 | cgtgagagaagcgcacacag |
| Fluc_15 | tctggttgccgaaaataggg |
| Fluc_16 | aatggcaccacgctcagaat |
| Fluc_17 | agggtggtgaacatgccgaa |
| Fluc_18 | aaagccgcaaatcaggtagc |
| Fluc_19 | aagcggtacatcagcaccac |
| Fluc_20 | agcagggcagactgaatttt |
| Fluc_21 | gcgaagaagctgaacagggt |
| Fluc_22 | cgtacttgtcgatcagggtg |
| Fluc_23 | aatctcgtgcaggttagaca |
| Fluc_24 | ctggcagatgaaagcgcttg |
| Fluc_25 | taatcagaatggcgctggtt |
| Fluc_26 | gaagaatggcaccaccttgc |
| Fluc_27 | gacataatcatagggccgcg |
| Fluc_28 | ctcagggttattcacgtagc |
| Fluc_29 | cttgtcgatcagggcgtttg |
| Fluc_30 | agtaggcaatgtcgccagag |
| Fluc_31 | ccacgatgaagaagtgctcg |
| Fluc_32 | ttgatcagagacttcaggcg |
| Fluc_33 | aaatgttagggtgctgcagc |
| Fluc_34 | acatagtccacgatctcctt |
| Fluc_35 | ttcttagccttgatcaggat |

smFISH probes toward *SunTag*

| Sequence Name | Sequence |
|---|---|
| SunTag_1 | ccacttcgttctcaagatga |
| SunTag_2 | cccttttcagtctagctac |
| SunTag_3 | aatttttgctcagcaactcc |
| SunTag_4 | ttctttagtcgtgctacttc |
| SunTag_5 | tttcgagagtaactcctcac |
| SunTag_6 | ccacttcgttttcgagatga |
| SunTag_7 | acttcccttttttaagcgtg |
| SunTag_8 | tcttggatagtagctcttca |
| SunTag_9 | acctcgttctcaagatgata |
| SunTag_10 | cggaacccttcttcaaacgc |
| SunTag_11 | agttcttcgagagcagttcc |
| SunTag_12 | gatcccttttttaatcgagc |
| SunTag_13 | tgaaagtagttcctcaccac |
| SunTag_14 | cttcgttttcgaggtggtaa |
| SunTag_15 | ccctgaacctttctttaatc |
| SunTag_16 | tactcagtaattcttcaccc |
| SunTag_17 | tttcgatagcaactcttcgc |
| SunTag_18 | tttttgagcctagcaacttc |
| SunTag_19 | ttttcgagagcaactcctcg |
| SunTag_20 | acctcattttccaagtggta |
| SunTag_21 | tttgctcaataactcctcgc |
| SunTag_22 | cgcgacttcgttctctaaat |
| SunTag_23 | ttcgataagagttcttcgcc |
| SunTag_24 | ctcattttcgaggtggtagt |
| SunTag_25 | agtggtagttcttgctcaag |
| SunTag_26 | ttcaatctcgcgacctcatt |
| SunTag_27 | attcttgctgagcaattcct |
| SunTag_28 | cgacttcgttctccaaatga |
| SunTag_29 | cgacttcattttccaagtgg |
| SunTag_30 | ttgctcaataactcttcgcc |
| SunTag_31 | ttcgttctccaagtggtaat |
| SunTag_32 | agttcttcgataagagctcc |
| SunTag_33 | gcgacttcattctctaagtg |
| SunTag_34 | ttcttgctcaagagctcttc |
| SunTag_35 | cacctcattttccaagtggt |
| SunTag_36 | ttagatagtaactcttcccc |
| SunTag_37 | cctcgttctcgagatgataa |
| SunTag_38 | gatagttcttcgacaggagt |
| SunTag_39 | ccttttttaagtcttgcaacc |
| SunTag_40 | ttactgagtagttcctcacc |
| SunTag_41 | ttcgttttccaggtggtaat |
| SunTag_42 | tcctgatcctttcttcaaac |
| SunTag_43 | cttttgagagcagttcttcg |
| SunTag_44 | gcaacctcattttccaaatg |
| SunTag_45 | tgccacttcccttttttaaa |
| SunTag_46 | tttcgacagaagttcctcac |
| SunTag_47 | gctacttcattctcgagatg |
| SunTag_48 | gagccagaacccttttttaag |
